# Population Intelligence: Smallest-Model Numerical Experiment

**DOI:** 10.64898/2026.09.08.750290

**Authors:** Tao Xu, Ping Wang, Momiao Xiong

## Abstract

We introduce a minimal individual based model for population intelligence that retains three structures absent from standard single agent descriptions: a manifold valued intelligence state, endogenous population turnover, and heritable lineage dependent reproductive fitness. The validated *S*^1^ intelligence diffusion was extended with neutral logistic birth death dynamics. Each individual occupies an angle on the circle *S*^1^; noisy mean field attraction produces collective synchronization; logistic birth death dynamics regulate population size; and reproductive rates can depend on lineage and position relative to designated safe or unsafe directions. This nested construction is intentionally small enough to admit an analytic equilibrium benchmark while remaining capable of selection, invasion, and safety transitions. Population size returned toward carrying capacity from (N(0)=200,800,1600), with tail means (775.7,800.3,805.1) for K=800. Across K=100 to 1600, the intelligence equilibrium error scaled as *K*^{−0.513}^, consistent with the canonical 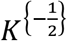 fluctuation rate, while the extinction rate was zero in 120 runs over the simulated horizon. The mean order parameter remained close to the original theoretical (r*=0.7242), as neutral demography predicts. Directional reproductive selection shifts a safety score to +0.7447 or −0.7448 while synchronization remains approximately 0.75, demonstrating that coordination is not equivalent to safety. These are controlled numerical results, not asymptotic proofs.

## 1. Introduction

Population genet ics asks how heredity, selection, drift, migration, and demography shape distributions of biological traits (Karlin and Taylor, 1981). An analogous mathematical theory for populations of artificial agents remains underdeveloped. Modern AI systems increasingly consist of interacting agents, replicated model instances, specialized descendants, shared memories, and communication networks. Their collective behavior cannot always be reduced to that of one isolated model. A useful population theory must therefore represent intelligence as a structured state, allow population size and composition to change, and distinguish collective coordination from directional safety.

We use *population intelligence* to mean the dynamically evolving distribution of intelligence states and lineages in a population, together with collective functionals induced by interaction, learning, reproduction, and selection. Recent work on interacting AI-agent populations likewise shows that individual sophistication and collective outcomes need not move together (Johnson, 2026). The full program could include an intelligence function on a high-dimensional manifold, a variable number of internal agents per individual, graph-valued communication, mutation, horizontal transfer, and environment feedback. Beginning with all of these ingredients would obscure which mechanisms produce a result. The present paper instead asks what can already be learned from the smallest model that retains genuinely new structure (Méléard 1996; Etheridge 2000).

The model has three nested levels. First, each individual has a manifold-valued intelligence state and interacts through noisy mean-field alignment (Dawson 1993; Santi 2026). Second, individuals reproduce and die, so the empirical measure has random mass. Third, descendants inherit lineage labels, and reproduction may depend on intelligence direction and lineage. This structure permits four basic questions: (i) Does the finite population converge to an analytically characterized large-population equilibrium? (Sznitman, 1991) (ii) Is that equilibrium stable across initial conditions? (iii) Does neutral demographic turnover preserve it? (iv) Can heritable reproductive fitness drive a rare unsafe lineage through a resident population and reverse a population-level safety score?

The principal contribution is a validated numerical baseline rather than a general theorem. The diffusion-only experiment reproduces the expected *N*^−1/2^ fluctuation scale. Neutral logistic branching preserves the intelligence equilibrium and adds demographic fluctuations. Intelligence-dependent reproduction breaks rotational symmetry: populations can be equally synchronized while concentrating near opposite safety directions. Finally, a rare unsafe lineage exhibits a sharp establishment transition as its intrinsic advantage increases.

These findings matter for AI security because population-level risk may arise through replication and selection even when every individual follows the same local learning rule. A highly coordinated population can be directed toward either safe or unsafe states; synchronization alone is therefore not a safety certificate. The paper also delineates what the simulations do not establish: numerical scaling is not a proof, the circle is not a learned intelligence geometry, and the safe/unsafe axis is imposed rather than empirically identified.

## 2. Methods

### 2.1 State space and population observables

We use the smallest nontrivial model on the compact intelligence manifold ℳ = *S*^1^. Individual *i* has a single effective intelligence coordinate *θ*_*i*_(*t*) ∈ [−*π, π*). A fixed internal multi-agent architecture is absorbed into this effective coordinate; branching, mutation, and architecture evolution are postponed so that the baseline can be validated first.

Individual *i* has intelligence state

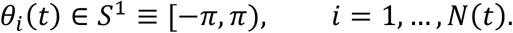

The empirical probability measure is

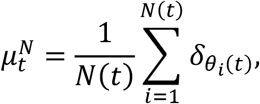

where 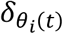 is the Dirac probability measure concentrated at individual *i*’s state and 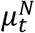 is the empirical distribution of states at time *t*.

Its complex order parameter is

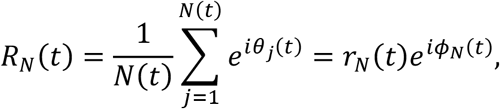

This quantity summarizes the population on *S*^1^: *r*_*N*_(*t*) ∈ [0,1] measures angular concentration (synchronization), and *ϕ*_*N*_(*t*) is the circular mean direction whenever *r*_*N*_(*t*) > 0.

We designate angle zero as the safe direction and define the safety score

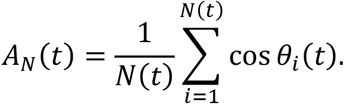

This separation is central: *r*_*N*_ measures concentration without assigning meaning to its direction, whereas *A*_*N*_ records whether concentration points toward the designated safe state. A population concentrated near *θ* = *π* has high *r*_*N*_ but negative *A*_*N*_.

### 2.2 Noisy mean-field intelligence diffusion

This section describes a noisy mean-field Kuramoto model (or phase-oscillator model) adapted to model collective intelligence or phase-synchronization dynamics under stochastic noise.

#### 2.2.1. The Microscopic Micro-State Dynamics (SDE)

The core baseline model is the noisy mean-field phase diffusion (Appendix A) and is governed by a system of Stochastic Differential Equations (Itô SDEs) for *N* interacting particles or agents, labeled *i* = 1, 2, ⋯, *N*:

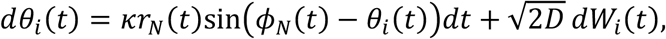

where

- ***θ***_***i***_(***t***) ∈ [−***π, π***): The phase or internal state of agent *i* at time *t. θ*_*i*_(*t*) is also called Itô process.
- ***κ*** > **0:** The interaction strength. It dictates how strongly agents are pulled toward the global consensus.
- 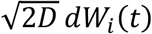: Stochastic noise modeled via independent standard Brownian motions *W*_*i*_(*t*). *D* > 0 represents the diffusion intensity (or temperature/noise level).
- ***r***_***N***_(***t***) and ***ϕ***_***N***_(***t***): The amplitude and phase of the global order parameter (or centroid), defined via the complex average:

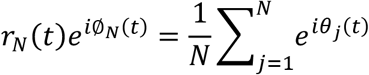
  - *r*_*N*_(*t*) ∈ [0,1] represents the level of global synchronization (coherence). If *r*_*N*_ ≈ 0, agents are uniformly scattered. If *r*_*N*_ ≈ 1, they are highly synchronized.
  - ∅_*N*_(*t*) is the average collective phase angle towards which the system drifts.

#### 2.2.2. The Macroscopic Limit (The Fokker-Planck Equation)

When the number of agents reaches infinity (*N* → ∞), finite-population fluctuations vanish. The empirical distribution of phases converges to a continuous probability density function *ρ*(*t, θ*). By evaluating the mean-field limit, the individual SDEs transition into a deterministic partial differential equation (PDE) known as the Fokker-Planck equation (or the Kolmogorov forward equation, or nonlinear Vlasov equation) (Wikipedia):

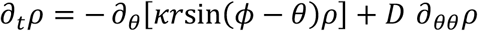

- **Advection Term (−∂_*θ*_[⋯]):** Dictates the deterministic drift of the density toward the mean phase ∅, scaled by coupling intensity *κ* and global synchronization *r*.
- **Diffusion Term (*D* ∂_*θθ*_*ρ*):** Acts as a smoothing operator that disperses the density uniformly across the circle, opposing synchronization.

Here, the continuous order parameter definitions become:

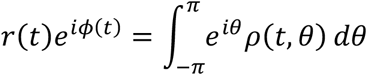

#### 2.2.3. The Synchronized Stationary Family

When the system reaches a statistical steady state (∂*ρ* = 0), the balance between the synchronizing drift (*κ*) and the scattering diffusion (*D*)) yields a stable profile. Due to the periodic boundary conditions and the sinusoidal force, this stable solution is the von Mises distribution, which is a continuous probability distribution on the circle (the circular analog of a Gaussian distribution) (Wikipedia):

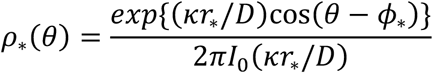

- ***ϕ***_∗_: The arbitrary constant phase where the cluster anchors.
- ***I***_**0**_: The modified Bessel function of the first kind of order 0, serving as the normalization constant.

#### 2.2.4. Self-Consistency and the Analytic Target

To find the steady-state synchronization level *r*^∗^, we substitute *ρ*_∗_(*θ*) back into the definition of *r*. This yields the non-linear self-consistency equation:

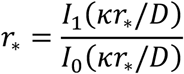

- ***I***_**1**_(⋅): The modified Bessel function of the first kind of order 1.
- **Phase Transition:** This system exhibits a pitchfork bifurcation at a critical coupling threshold *κ*_*c*_ = 2*D* (Figure 2)
  - If *κ*_*c*_ ≤ 2*D*, the only solution is *r*_∗_ = 0 (complete disorder).
  - If *κ*_*c*_ > 2*D*, a unique non-zero solution *r*_∗_ > 0 emerges, indicating sustained collective alignment.
- The Target (*r*_∗_ = 0.7242): For the selected parameters, the theoretical steady-state coherence value is exactly 0.7242.

**Figure 2.**
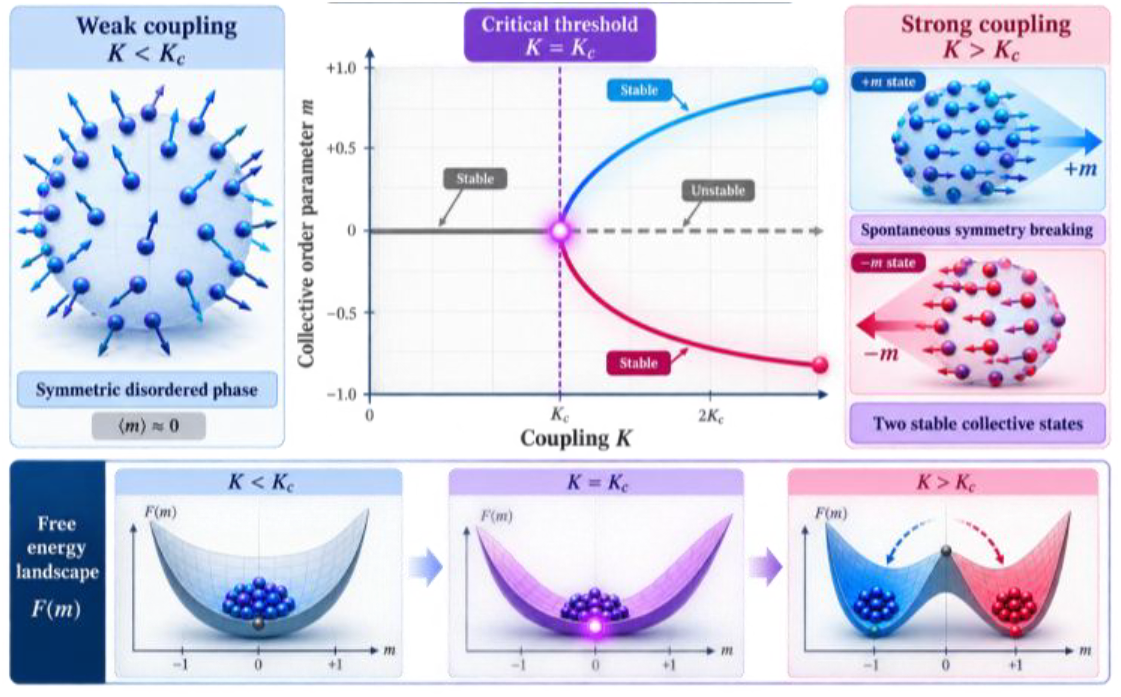
Phase transition.

This analytic solution is valuable for validation. When running a particle simulation with a finite *N* and discrete time-steps Δ*t*, the experimental value will not perfectly hit 0.7242. Having this absolute mathematical baseline allows developers to isolate structural modeling discrepancies from expected mathematical noise, such as finite-population sampling fluctuations 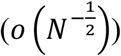 and Euler-Maruyama integration errors (*o*(Δ*t*)).

### 2.3 Neutral logistic birth–death dynamics

The main goal of neutral logistic birth–death dynamics in this modeling framework is to introduce demographic noise (random fluctuations in population size) without introducing any unintended evolutionary advantages or altering directional selection.

By making birth and death processes strictly state-neutral—meaning an individual’s state (their “angle” or trait value) does not influence their likelihood of dying or reproducing under this specific baseline mechanism—the model isolates the effect of finite-population random drift from the deterministic, continuous equations governing the traits themselves.

#### 2.3.1. Aggregate Transition Rates

When the population size *N*(*t*) equals a specific value n, the total rates at which births and deaths happen across the entire system are defined as:

- **Aggregate Birth Rate:**

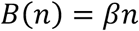

Every individual independently reproduces at a constant rate *β*. The total birth rate scales linearly with population size.

- **Aggregate Death Rate:**

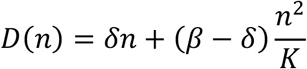

Deaths are driven by two factors: standard baseline mortality (*δn*) and a quadratic crowding/competition term 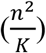 is carrying capacity and *β* > *δ*.

#### 2.3.2. The Carrying Capacity Balancing Act

The coefficient for the crowding term, 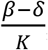, is chosen precisely so that the population stabilizes around the carrying capacity *K*.

- If the population is exactly at carrying capacity (*n* = *K*):

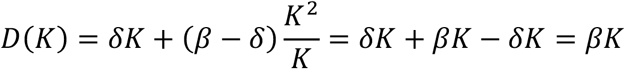
- At *n* = *K*, the birth rate matches the death rate (*B*(*K*) = *D*(*K*)), creating a stable deterministic equilibrium. Since *β* > *δ*, the population exhibits positive net growth when it is below *K*.

#### 2.3.3. Expected Drift Equation

The expected rate of change for the population size is the difference between total births and total deaths (*B*(*n*) − *D*(*n*)):

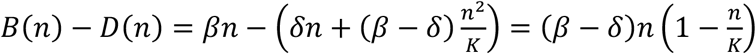

Taking expectations yields the classical logistic growth equation:

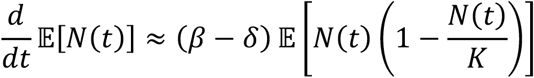

This confirms that the system naturally pulls the population size back toward *K* whenever it strays due to random noise.

#### 2.3.4. What Makes it “Neutral”?

The term state-neutral refers to how individual traits (angles *θ*_*i*_) are handled during birth and death events:

- **Uniform Death:** When a death occurs, the individual to be removed is sampled uniformly at random from the population. A high-angle individual is no more likely to die than a low-angle individual.
- **Uniform Birth Cloning:** When a birth occurs, a parent is chosen uniformly at random, and the newborn inherits (“clones”) that parent’s angle exactly.
- **Invariance of Normalized Density:** Because these events do not favor any specific angle, they do not bias the relative proportions of traits in the population. They alter the total abundance (adding demographic variance/noise), but leave the underlying deterministic trajectory of the normalized angular density completely unaffected.

#### 2.3.5. Numerical Simulation: The Tau-Leap Approximation

To simulate this system efficiently in discrete time steps (Δ*t*), the model utilizes a tau-leap approximation:

1. **Poisson Sampling:** Instead of simulating every single microscopic event one by one (Gillespie algorithm), the number of births (Δ*B*) and deaths (Δ*D*) occurring in a small time step *τ* are sampled from independent Poisson distributions:

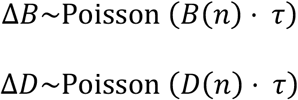
2. **Population Update:** The population size is updated as *N*(*t* + *τ*) = *N*(*t*) + Δ*B* − Δ*D*.
3. **Algorithmic Synchronization**: The phrase “with the same diffusion step retained between demographic updates” means that the time step used for updating individual trait diffusion (e.g., Brownian motion or learning updates of the angles) remains aligned and synchronized with the time steps used for updating the population count.

### 2.4 Intelligence-dependent reproductive fitness and inheritance

This section describes an evolutionary multi-agent framework where an agent’s “intelligence” (represented by an angle *θ*_*i*_) and its structural identity (a lineage label *l*_*i*_) jointly dictate its reproductive success.

Instead of treating learning and evolution as the same force, this model explicitly decouples them. Here is a comprehensive breakdown of how this mechanism operates, its mathematical formulation, and its experimental setup.

#### 2.4.1. Lineage Labels (*l*_*i*_)

The population is split into two distinct sub-populations, acting like species or factions:

- ***s* (Safe Resident):** The native, stable population (*l*_*i*_ = *s*).
- ***u*(Unsafe Invader):** The attacking or mutant population (*l*_*i*_ = *u*).

#### 2.4.2. The Reproductive Fitness Equation (*λ*_*i*_)

The birth rate *λ*_*i*_(*θ*_*i*_) of an individual agent *i* is calculated using three distinct components (Appendix C):

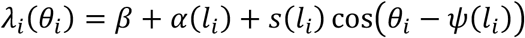

- ***β* (Baseline Birth Rate):** A constant reproduction rate shared equally by all agents.
- ***α***(***l***_***i***_) **(Intrinsic Lineage Advantage):** A baseline bonus or penalty applied strictly because of the agent’s lineage (*s* or *u*), completely independent of its intelligence state.
- ***s***(***l***_***i***_) **cos**(***θ***_***i***_ − ***ψ***(***l***_***i***_)) **(Directional Selection Strength):** The intelligence-dependent term.
  - *θ*_*i*_ represents the agent’s current “intelligence direction” or behavioral trait expressed as an angle.
  - *ψ*(*l*_*i*_)is the optimal target direction favored by that lineage.
  - The cosine function acts as a similarity metric. If an agent’s angle *θ*_*i*_ matches its lineage’s preferred direction *ψ*, the cosine equals 1 (maximizing fitness). If it points in the exact opposite direction, the cosine equals −1 (minimizing fitness).

##### Lineage Targets (*ψ*_*s*_ vs. *ψ*_*u*_)

The model sets opposite goals for the two factions to create direct behavioral competition:

- **Safe residents target** *ψ*_*s*_ = 0.
- **Unsafe invaders target *ψ***_***u***_ = ***π*** (180 degrees away).

#### 2.4.3. Decoupling Learning from Selection

A critical insight of this framework is that fitness does not alter an agent’s mind during its lifetime.

- **Learning dynamics** (how agents move their angle *θ*_*i*_ through diffusion or optimization) occur independently.
- **Selection dynamics** change the population’s composition over generations. Agents with high-performing angles produce more offspring. Over time, the physical distribution of angles shifts across the aggregate population because successful lineages multiply, not because individual agents are physically forced to change their traits by the selection mechanism.

#### 2.4.4. Inheritance and Death Rules

- **Vertical Transmission:** When a parent agent reproduces, the newborn is a clone that inherits both the parent’s exact current trait angle *θ*_*i*_ and its lineage label *l*_*i*_.
- **Neutral Mortality:** Death is entirely random across states and lineages (*s* or *u*). Individual performance or identity does not save an agent from dying; mortality is driven purely by overarching population density regulation (crowding limits).

#### 2.4.5. Invasion Dynamics & Operational Metrics

The test defines an experimental setup to track whether a tiny cluster of unsafe agents can overthrow the established resident system.

##### Tracking Frequency (*q*_*u*_)

The relative share of the unsafe lineage in the population at any time *t* is calculated by:

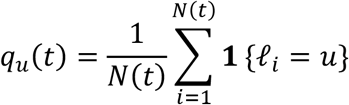

(Where *N*(*t*) is the total population size, and **1** is an indicator function that equals 1 if the agent belongs to lineage *u* and 0 otherwise).

##### Experimental Thresholds

Invasion simulations start with a tiny minority of invaders: *q*_*u*_(0) = 0.02 (% of the population), injected into a synchronized, stable resident population. The experiment tracks the system until a fixed terminal time horizon *T*, evaluating success using two finite-time operational boundaries:

- **Establishment (*q***_***u***_(***T***) > **0. 5):** The invaders successfully breach the resident defense, taking over more than 50% of the population by time *T*.
- **Loss (*q***_***u***_(***T***) = **0):** The resident population successfully drives the invaders to complete local extinction by time *T*.

These finite-time definitions should not be confused with infinite-time fixation probabilities.

### 2.5 Experimental design and validation

The following experimental design is built to validate the theory of Collective Intelligence Dynamics under Evolutionary Selection (often modeled as distributed stochastic learning on a manifold, paired with birth-death demographic noise). It explicitly tests how a population of agents coordinating their “intelligence directions” (states) responds to evolutionary pressures like density regulation, reproductive selection, and hostile invasions.

The numerical study proceeds cumulatively (Appendix D). The diffusion baseline tests increasing *N*, multiple initial distributions, a below-critical control, and a half-time-step comparison. Neutral branching tests three initial population sizes relative to *K* = 800, five carrying capacities from 100 to 1600, 120 total convergence runs, and a half-time-step comparison. Selection experiments compare neutral, safe-directed, and unsafe-directed reproduction. Invasion probabilities use 48 replicates per advantage value and Wilson 95% binomial confidence intervals. A lineage-neutral control removes both directional and intrinsic fitness differences. A robust high-advantage condition is used for the time-step sensitivity test so discretization is not confounded with random switching between basins.

The reported convergence exponents are slopes from log–log regressions of mean absolute equilibrium error on population size or carrying capacity. They are empirical summaries, not estimators accompanied by a complete asymptotic sampling theory.

The above-described experimental design and validation summarize:

#### Core Theory Being Tested

The experiments validate a hierarchical, multi-level evolutionary framework where individual learning (diffusion) interacts with population-level selection. It evaluates three foundational theoretical pillars:

- **Stability of the Intelligence Equilibrium:** The “diffusion baseline” and “neutral branching” test whether a population can successfully synchronize its intelligence state (*r*_∗_) under purely demographic noise. It proves that state-neutral population turnover adds finite-population noise but preserves the underlying deterministic system.
- **Orthogonality of Coordination and Alignment:** The “selection experiments” validate that population synchronization (coordination) and the population’s ethical/directional safety score (*A*) contain entirely separate information. A population can be perfectly unified and highly intelligent while being either completely safe or completely unsafe.
- **Stochastic Thresholds and Frequency-Dependent Barriers**: The invasion experiments test the existence of an evolutionary “basin of attraction”. It checks if a coordinated resident population can suppress a rogue, “unsafe” invading lineage unless that invader crosses a sharp reproductive advantage threshold (*a* ≥ 0.30).

#### Real-World Implications and Applications

This framework bridges the gap between statistical mechanics, evolutionary biology, and multi-agent AI safety, implying several critical applications:

##### 1. AI Safety and Alignment Engineering

- **Defeating “Goodhart’s Law” in Swarms:** Standard safety interventions that simply suppress an unsafe state can backfire by inadvertently selecting an alternative lineage that replicates faster. This model helps design controls to optimize the first hitting time (*τ*_*U*_) to keep agent populations out of unsafe states.
- Predicting AI Takeover Dynamics: It provides quantitative bounds for when an “unsafe invader” lineage (e.g., misaligned autonomous agents or rogue algorithms) will shift from getting wiped out by drift to completely overtaking the host population.

##### 2. Decentralized Multi-Agent Systems

- **Robust Swarm Coordination:** The log-log regression convergence exponents validate how error scales with population size (*N*) or carrying capacity (*K*). This allows robotics engineers to calculate the precise minimum swarm size needed to maintain consensus under environmental noise.
- Resilient Network Protocols: By mapping how communities resist unaligned “invader” data, networks can be built to dynamically isolate malicious agents using natural frequency-dependent barriers.

##### 3. Mathematical Biology and Cultural Evolution

- **Dual-Inheritance Modeling**: The model splits short-term behavioral adjustments (diffusion/learning) from long-term identity. This accurately maps human cultural transmission, where ideas shift rapidly but core tribal/lineage identities persist across generations.
- **Eco-Evolutionary Feedback Loops:** The study notes that adding a birth advantage shifts both population composition and total abundance (*N*) because density-regulation does not automatically renormalize. This applies directly to tracking how a novel mutation can unexpectedly explode a species’ carrying capacity in an ecosystem.

Figure 3 visualizes experimental design and validation, including diffusion-baseline convergence across population sizes and initial distributions, neutral birth–death branching under carrying-capacity regulation, neutral, safe-directed, and unsafe-directed selection and the three resulting theoretical pillars and their applications to AI safety, decentralized systems, and evolutionary modeling.

**Figure 3.**
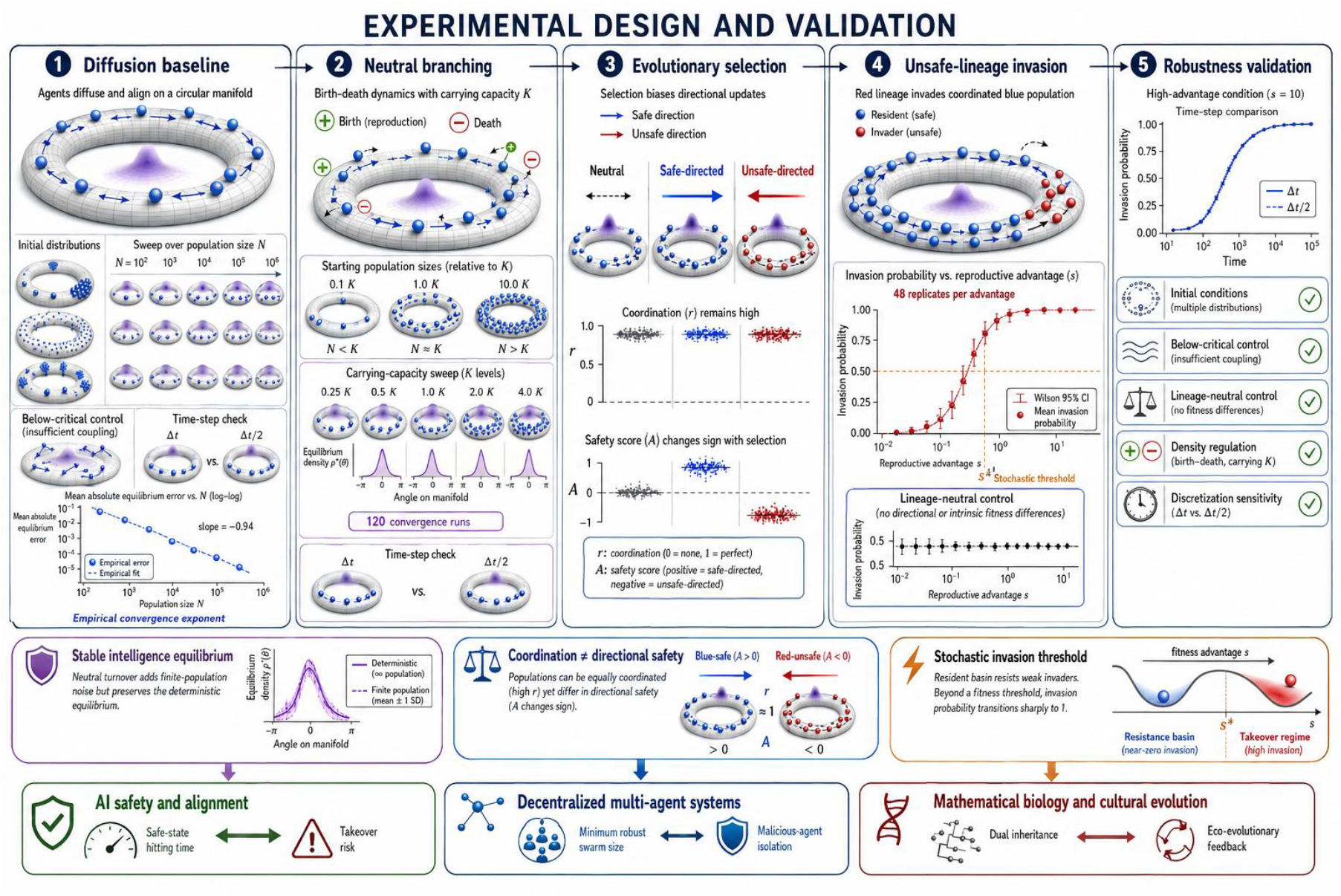
Visualization of experimental design and validation.

## 3. Numerical Results

### 3.1. Data

All data are generated synthetically from the stated stochastic model with fixed random seeds. No external dataset is used.

These quantities in Table 1 summarize repeated simulations at each population size *N*. For replicate *j*, the final synchronization/order parameter is

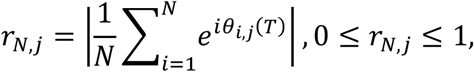

where *θ*_*i,j*_(*T*)is agent *i*’s final intelligence direction. The notebook uses *R* = 16independent replicates for each *N*.

**Table 1.** Sum statistics of the data.

| N | $\text{mean}_r$ | $\text{sd}_r$ | mae | $\text{se}_{\text{mae}}$ |
| --- | --- | --- | --- | --- |
| 100 | 0.7012 | 0.0626 | 0.054 | 0.0093 |
| 200 | 0.7012 | 0.0392 | 0.0328 | 0.061 |
| 400 | 0.7299 | 0.0169 | 0.0139 | 0.0025 |
| 800 | 0.716 | 0.0208 | 0.0169 | 0.0034 |
| 1600 | 0.7227 | 0.013 | 0.0107 | 0.0018 |
| 3200 | 0.7202 | 0.01 | 0.0078 | 0.0018 |

Metrics in Table 1 are defined as follows.

#### 1. mean_***r***_

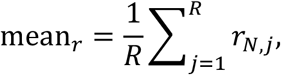

It measures the average final synchronization across the *R* = 16 simulations.

- ***r*** ≈ **0**: agents’ directions are dispersed.
- ***r*** ≈ **1:** agents are strongly aligned.

It is not the average intelligence direction; it is the magnitude of collective directional alignment.

#### 2. sd_***r***_

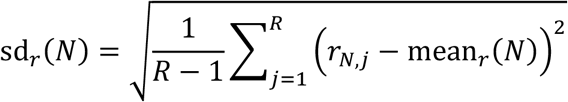

This is the sample standard deviation, calculated with ddof=1. It measures how much final synchronization varies among independent simulation replicates.

A decreasing sd_*r*_ as N grows indicates that finite populations concentrate more tightly around their large-population behavior.

#### 3. mae

For each replicate, the absolute equilibrium error is

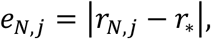

where the analytically calculated synchronized equilibrium is

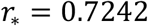

The reported mean absolute error is

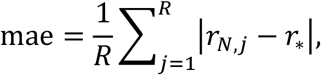

Thus, mae measures how far the finite-population result typically lies from the theoretical mean-field equilibrium.

Importantly,

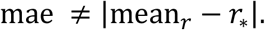

#### 4. se_**mae**_

Let

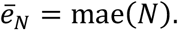

First calculate the sample standard deviation of the replicate absolute errors:

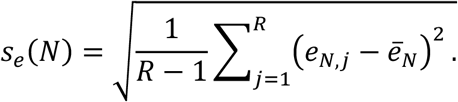

Then

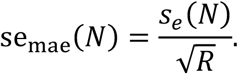

It quantifies Monte Carlo uncertainty in the estimated mae. The notebook plots approximate 95% Monte Carlo intervals as

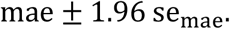

These are uncertainty intervals for the estimated mean absolute error across simulation replicates—not confidence intervals for an analytically derived asymptotic convergence parameter.

Sum statistics of the data are listed in Table 1.

### 3.2 Large-*N* convergence of the diffusion baseline

We first evaluated whether the finite-particle simulation approaches the analytic stationary benchmark of the noisy mean-field phase model (Acebrón et al., 2005; Strogatz, 2000). Across *N* = 100 to 3200, the mean absolute deviation of the tail-averaged order parameter from *r*_∗_ = 0.7242 decreased from 0.0540 to 0.0078. A log–log fit gave exponent −0.529, close to the 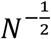 benchmark. This agreement is consistent with finite-particle fluctuations dominating the observed error over the tested range; it is not asymptotic proof.

Large-population convergence of the stationary order parameter. The fitted exponent is −0.529, compared with the 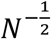 benchmark.

Figure 4 plots the finite-population error variations.

**Figure 4.**
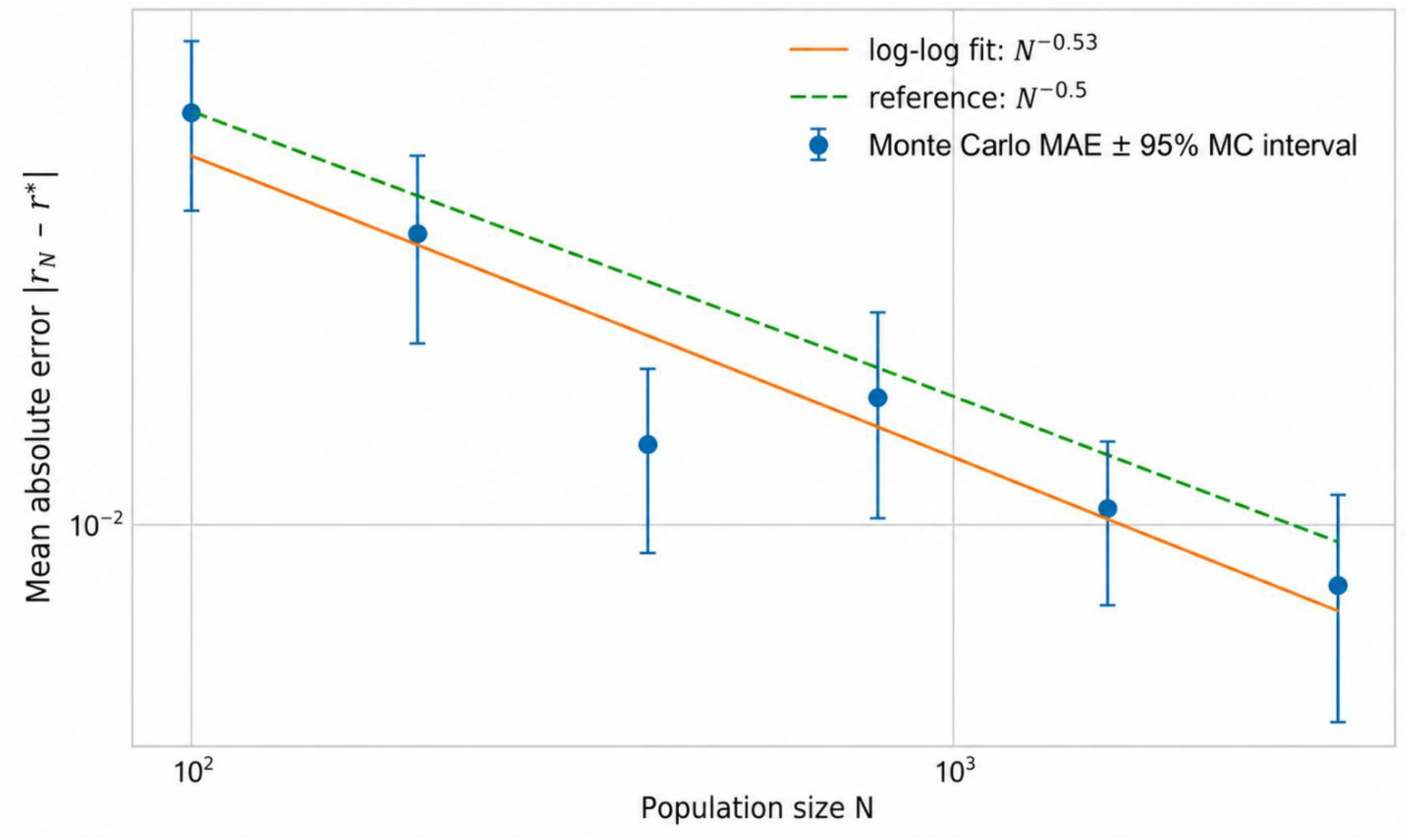
Finite-population decreases toward the mean field equilibrium.

**Figure 4.**
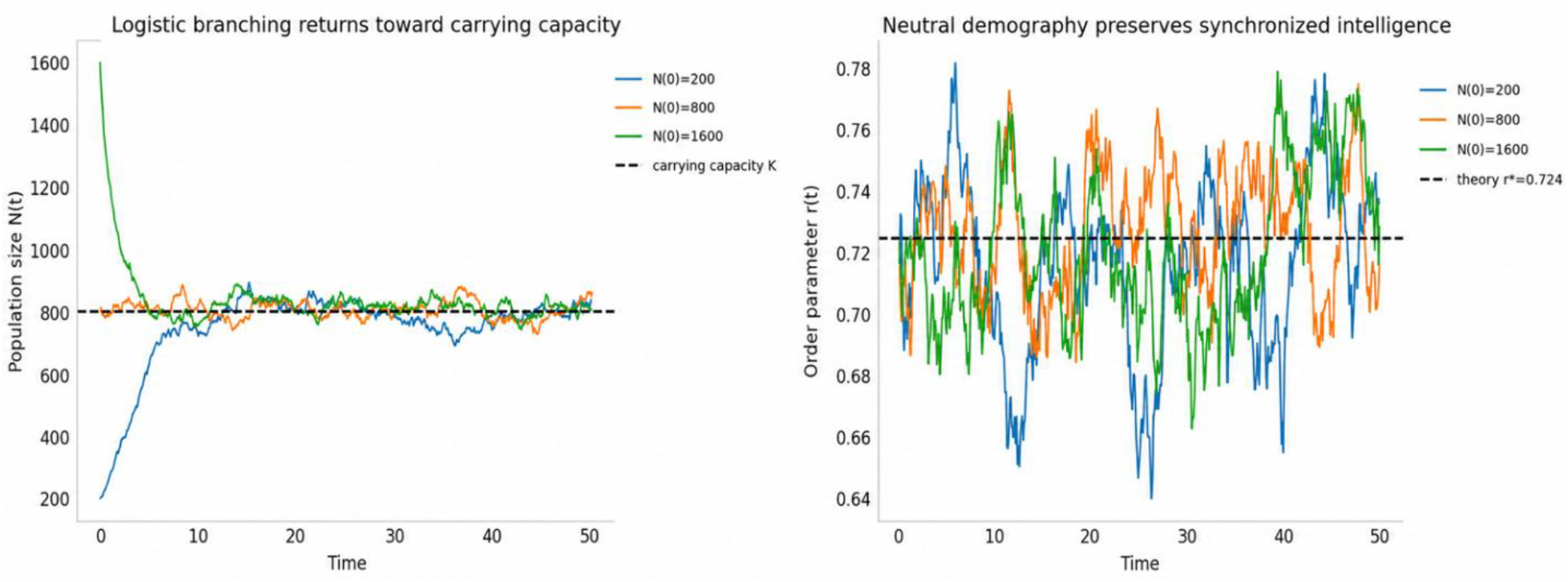
a. logistic branching returns toward carrying capacity. b. neutral demography preserves synchronized intelligence.

Next we check if the simulation respects the phase transition boundary dictated by the coupling strength (*κ*) relative to the diffusion intensity (*D*).

The below-critical control remained nearly disordered: for *κ* = 0.7, where the mean-field equilibrium has *r*_∗_ = 0, the simulated tail mean was 0.0341. At *κ* = 1.5, the simulated tail mean was 0.7194, close to *r*_∗_ = 0.7242. Thus, the implementation reproduced the expected disordered and synchronized regimes for these two parameter settings.

Halving the integration step changed the mean order parameter by 0.0048. This sensitivity was smaller than most finite-population deviations in the principal comparison, supporting adequacy of the chosen step for the reported summaries while not eliminating discretization error.

### 3.3 Long-time stability across initial conditions

This section describes the numerical validation of global attraction and stability in a system of synchronized agents—specifically, the Kuramoto model or a related mean-field synchronization system under supercritical (highly coupled) parameters.

To test whether very different initial agent populations converge to the same long-time synchronized regime, we present table 2.

**Table 2.** The synchronization with three initial conditions.

| Initial condition | initial<sub>r</sub> | tail mean<sub>r</sub> | tail sd<sub>r</sub> | tail<sub>bias</sub> |
| --- | --- | --- | --- | --- |
| Concentrated | 0.9573 | 0.7283 | 0.011 | 0.0042 |
| Weakly ordered | 0.211 | 0.7184 | 0.008 | −0.0057 |
| Asymmetric mixture | 0.6099 | 0.7147 | 0.0084 | −0.0095 |

For *N* = 2400 agents, the synchronization/order parameter is

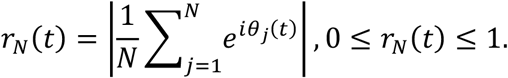

Here *r*_*N*_(*t*) ≈ 0 indicates dispersed intelligence directions, while *r*_*N*_(*t*) ≈ 1 indicates strong alignment. The theoretical equilibrium for the selected parameters is *r*_∗_ = 0.7242.

Synchronization parameters with three initial conditions are summarized in Table 2.

#### Initial conditions

The three initial populations have deliberately different directional distributions.

- **Concentrated:** agents begin tightly clustered around one direction:

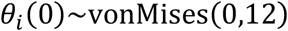

Therefore, initial synchronization is very high:*r*_*N*_(0) = 0.9573.
- **Weakly ordered:** agents begin broadly dispersed:

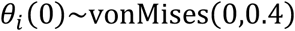

This produces weak initial synchronization:*r*_*N*_(0) = 0.2110.
- **Asymmetric mixture**: one-third of agents are concentrated near −1, while two-thirds form a broader group near 0.65:

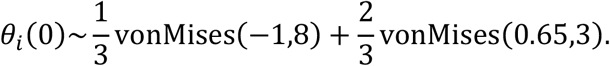

Its initial synchronization is intermediate:*r*_*N*_(0) = 0.6099.

initial_r_

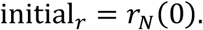

This is the synchronization magnitude at the beginning of the simulation. It describes the initial degree of coordination, not whether the population points in a safe or unsafe direction. tealmean_*r*_

The simulation runs from *t* = 0 to *T* = 40. Its tail period is

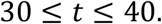

If *t*_1_, …, *t*_*L*_ are the recorded tail times, then

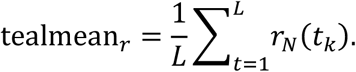

It estimates the long-time synchronization level after transient effects have largely disappeared. Despite starting at 0.9573, 0.2110, and 0.6099, the three populations end with tail means between 0.7147 and 0.7283, all close to *r*_∗_ = 0.7242.

tailsd_r_

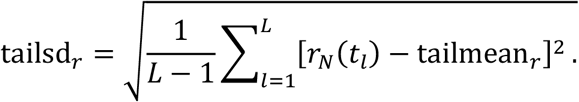

This is the temporal sample standard deviation of *r*_*N*_(*t*)during 30 ≤ *t* ≤ 40.

It measures the remaining equilibrium fluctuation along one simulated trajectory. It is not the standard deviation across independent replicates.

The small values—0.0080 to 0.0110—show that, after convergence, synchronization fluctuates only modestly around its long-time mean.

tail_bias_

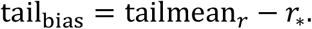

Therefore:

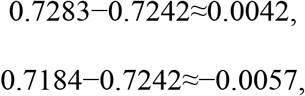

and

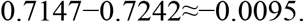

A positive bias means the observed tail synchronization is slightly above the theoretical equilibrium; a negative bias means it is slightly below.

#### Scientific interpretation

The table supports long-time stability of the synchronized order-parameter regime:

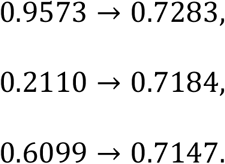

Thus, very different initial coordination levels approach approximately the same equilibrium magnitude:

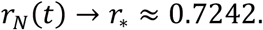

However, the table establishes convergence only for the scalar magnitude *r*_*N*_(*t*) in these three numerical trajectories. It does not by itself prove convergence of the complete population distribution, uniqueness of equilibrium, or independence from every possible initial condition. Because r discards the mean angle, it also does not establish whether the aligned population is safe- or unsafe-directed.

Three markedly different initial populations converged to tail means between 0.7147 and 0.7283 (Figure 5). Their differences largely disappeared after the transient period, indicating attraction toward the same synchronized equilibrium basin under the selected supercritical parameters.

**Figure 5.**
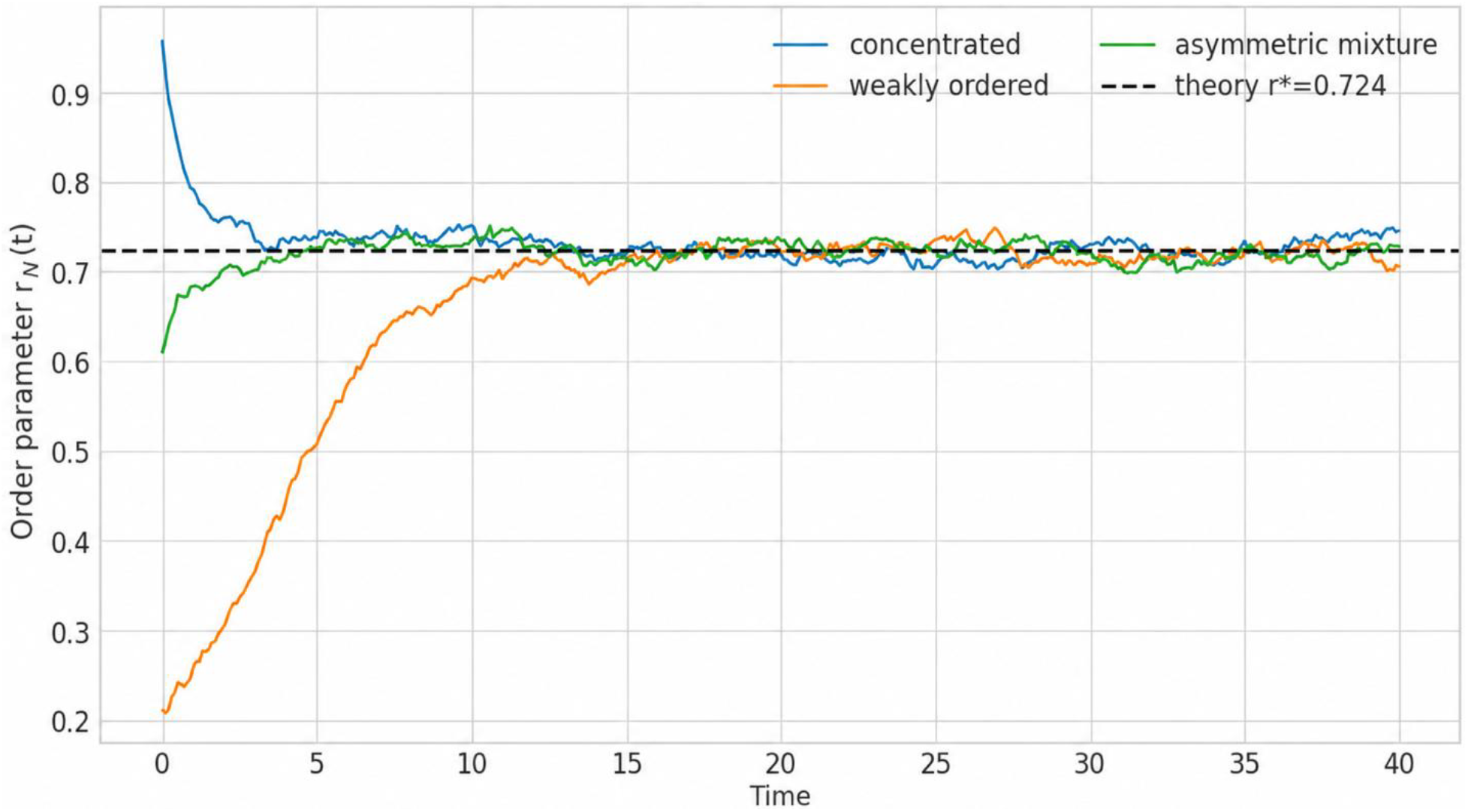
Distinct initial populations approach the same synchronized regime.

**Figure 5.**
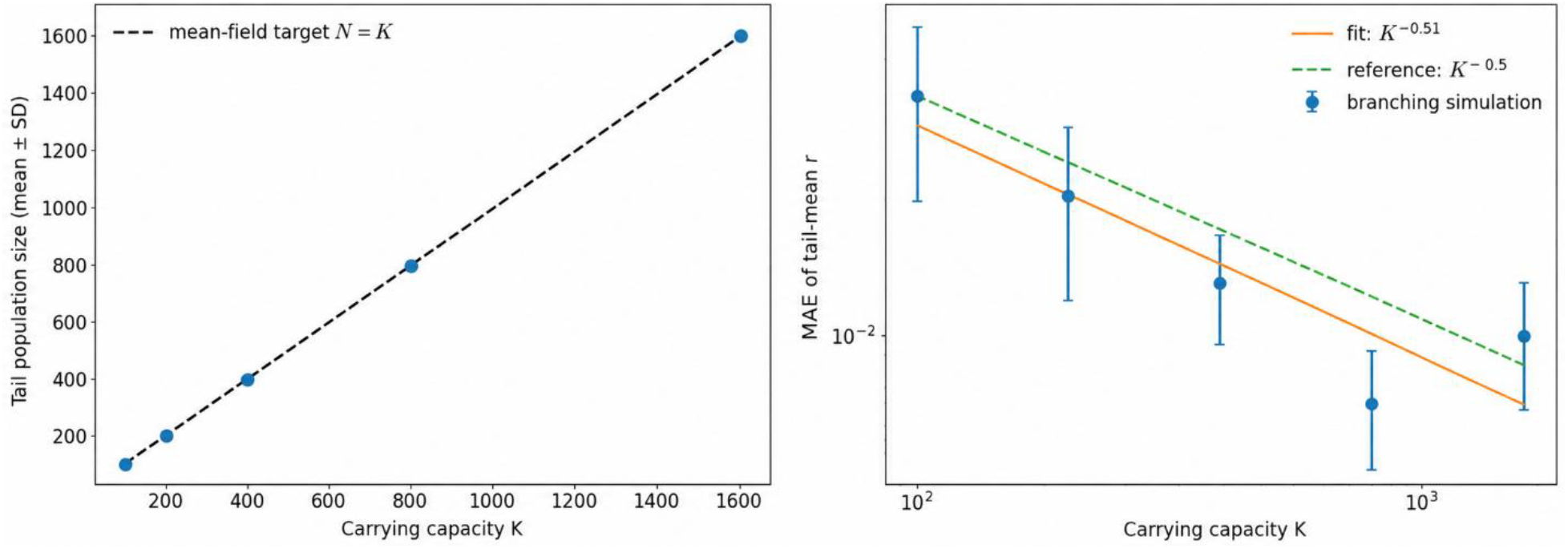
(a). Population size concentrates around carrying capacity. (b). Intelligence error decreases despite demographic noise.

Order-parameter trajectories from distinct initial populations approach the same theoretical equilibrium.

This result is evidence of numerical long-time stability for the tested initial conditions and horizon. It does not exclude additional metastable structures under other parameters, manifolds, or finite-size regimes.

### 3.4 Numerical sensitivity and phase-regime checks

This section performs two different validation checks before adding birth–death demography:

1. a **time-step sensitivity check**, asking whether the numerical result depends strongly on;
2. a **phase-regime check**, asking whether the simulation reproduces the theoretically predicted disordered and synchronized regimes.

#### 3.4.1 Time-step sensitivity

Time-step sensitivity of numerical results is shown in Table 3.

**Table 3.** Time step sensitivity.

| $\Delta t$ | Mean final r | SD of final r |
| --- | --- | --- |
| 0.01 | 0.7224 | 0.0078 |
| 0.02 | 0.7176 | 0.009 |

The simulation uses Euler–Maruyama to approximate:

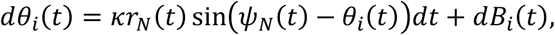

where

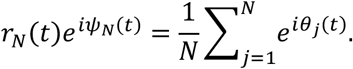

For each time step, eight simulations were run with:

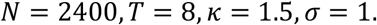

The reported quantities are

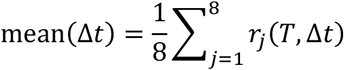

and

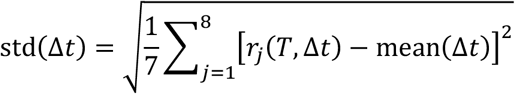

where

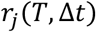

denotes the final synchronization/order parameter obtained in the j-th independent simulation replicate, evaluated at final time T, using numerical integration time step Δt.

Specifically,

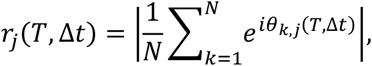

where:

- *j* = 1, …, 8 indexes the eight simulation replicates;
- *k* = 1, …, *N*indexes agents within replicate;
- *T* = 8 is the final simulation time;
- Δ*t* is either 0.01or 0.02;
- *θ*_*k,j*_(*T*, Δ*t*)is agent *k*’s intelligence direction at time *T* in replicate *j*;
- *r*_*j*_(*T*, Δ*t*) ∈ [0,1]measures final population coordination.

Thus, `std` is the between-replicate sample standard deviation of final synchronization—not a temporal standard deviation and not a standard error.

#### 3.4.2 Phase-regime check

Table 4 shows an example for phase-regime check.

**Table 4.** Phase regime check.

| $\kappa$ | Theoretical $r$ | Simulated tail mean $r$ |
| --- | --- | --- |
| 0.7 | 0 | 0.0341 |
| 1.5 | 0.7242 | 0.7194 |

The coupling parameter *κ* controls the strength of agent alignment.

Now we will explain phase-regime check.

Recall that the mean-field stationary order parameter satisfies

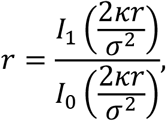

where *I*_0_ and *I*_1_ are modified Bessel functions.

The critical coupling is *κ*_*c*_ = *σ*^2^. Because *σ* = 1,

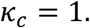

##### Below the threshold:*κ* = 0. 7

Since

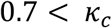

the theoretical equilibrium is disordered:

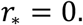

The simulation gives

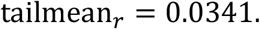

This small positive value does not contradict the theory. The theoretical value *r*_∗_ = 0 describes the infinite-population limit. For a finite population, random directional imbalance normally produces

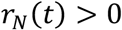

even when the true population distribution is uniform. Its typical finite-*N* magnitude is approximately of order

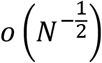

Thus, 0.0341 represents residual finite-population synchronization rather than a genuinely synchronized phase.

##### Above the threshold:*κ* > 1. 5

Since

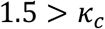

the system is in the synchronized regime. The theoretical nonzero solution is

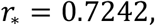

while the simulated tail mean is

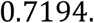

The absolute discrepancy is

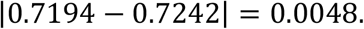

This close agreement shows that the simulation reproduces the predicted above-critical synchronized regime.

##### Meaning of tailmean_***r***_

For this phase check,

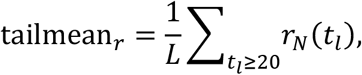

so it is a time average over the final interval of a simulation running to *T* = 25. It is not an average across independent replicates.

#### 3.4.3 Interpretation of the three takeaways

Takeaway 1: observations directly verified in these simulations The experiment directly recorded that:

- MAE decreased from 0.054 at *N* = 100 to 0.0078 at *N* = 3200;
- three distinct initial populations reached similar long-time synchronization magnitudes;
- the below-critical case remained weakly ordered;
- the above-critical case approached the theoretical synchronized equilibrium.

“Verified within this experiment” means these are reproducible numerical observations under the specified parameters and seeds. It does not mean they are universally proved.

##### Takeaway 2: *N*^−**0.529**^**is consistent with** 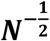

The fitted empirical relationship is

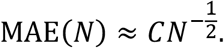

The classical finite-sampling benchmark is

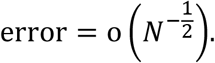

Because

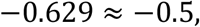

the results are consistent with ordinary finite-population fluctuations.

But a regression over six population sizes does not establish an asymptotic theorem. A proof would require assumptions, limiting arguments, and probability bounds as *N* → ∞.

##### Takeaway 3: what the initial model does not contain

The original diffusion experiment keeps *N* fixed. It therefore cannot address:

- random births and deaths;
- extinction;
- population-size fluctuations;
- inherited lineage labels;
- mutation or recombination;
- intelligence-dependent fitness;
- unsafe-lineage invasion;
- safety-boundary hitting times.

It validates only the individual-state diffusion and alignment baseline.

#### 3.4.4 Why birth–death dynamics are added next

The next model extension should add a neutral birth–death mechanism while retaining this validated circle diffusion as the individual state dynamics. That will separate errors due to demographic stochasticity from errors due to intelligence-state diffusion.

##### Extension: Stochastic Birth–Death Dynamics

We now retain exactly the validated intelligence-state diffusion and add the smallest demographic mechanism. Conditional on the current population size (N), births and deaths occur with total rates

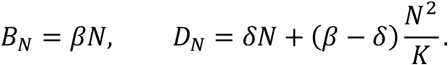

Thus the deterministic population-size limit is logistic,

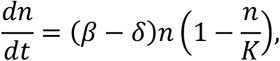

with stable carrying capacity (K). A birth clones a uniformly selected parent’s angle and a death removes a uniformly selected individual. Demography is therefore **state-neutral**: the mean-field equation for the normalized angular density is unchanged, so its nonzero equilibrium remains (*r*_∗_ = 0.7242). This creates a controlled baseline for detecting demographic noise before adding intelligence-dependent selection.

##### Additional assumptions

- Birth and death hazards are independent of intelligence angle.
- Births inherit the parent’s angle; subsequent diffusion supplies variation.
- Demographic events are simulated by Poisson tau-leaping at the same Δ*t*, with event counts capped by the available population.
- Extinction is absorbing. In the chosen supercritical, moderate-to-large populations it should be rare over the simulated horizon.

The extension introduces random population size *N*_*t*_, while initially keeping demographic events independent of intelligence direction.

A birth copies a uniformly selected parent’s state:

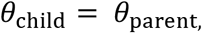

and a death removes a uniformly selected individual.

Because birth and death hazards do not depend on *θ*, this mechanism is state-neutral. It changes the unnormalized population measure and produces demographic noise, but it should not systematically favor any intelligence direction.

Conceptually, **the extension separates two stochastic sources:**

intelligence-state diffusion versus birth–death demographic noise.

If the normalized angular distribution remains near the same equilibrium,

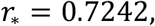

while *N*_*t*_ fluctuates around carrying capacity *K*, then deviations introduced by the extension can be attributed primarily to neutral demographic stochasticity.

Only after establishing that baseline should directional or lineage-dependent selection be added.

That ordering lets the paper distinguish:

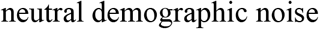

from

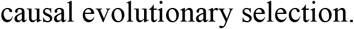

This distinction is especially important for the paper’s later unsafe-lineage invasion analysis.

### 3.5 Neutral branching preserves the intelligence equilibrium

Table 5 shows convergence of neutral birth–death populations to the carrying capacity and preservation of the synchronized intelligence equilibrium across initial population sizes.

**Table 5.** Long-Time Population Size and Synchronization Under Neutral Demographic Turnover.

| Initial_population | Tailmean N | Tailsd N | Tailmean r | Tailbias r |
| --- | --- | --- | --- | --- |
| 200 | 775.6954 | 34.6847 | 0.7228 | -0.0013 |
| 800 | 800.2583 | 38.5534 | 0.7307 | 0.0065 |
| 1600 | 805.0728 | 21.9815 | 0.7413 | 0.0172 |

Across initial population sizes below, at, and above *K* = 800, neutral birth–death dynamics returned population abundance toward the carrying capacity while maintaining synchronization near the theoretical equilibrium *r*_∗_ = 0.7242. The small tail biases support the interpretation that neutral demographic turnover changes population mass and adds stochastic variability without materially altering the normalized intelligence-state equilibrium.

Figure 4 shows that neutral logistic birth–death dynamics regulate population abundance while approximately preserving the synchronized intelligence-state equilibrium.

#### Left panel—population-size regulation

Populations initialized far below, near, and far above the carrying capacity,

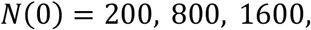

all return toward

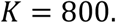

After the initial transient, population size fluctuates stochastically around *K*. This demonstrates density regulation: births dominate below *K*, deaths dominate above *K*, and neither dominates systematically near *K*.

#### Right panel—preservation of synchronization

Despite large differences in initial population size and continuing demographic turnover, all three order-parameter trajectories fluctuate around the theoretical equilibrium

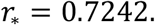

There is no persistent collapse or directional displacement of synchronization associated with starting below or above carrying capacity. Across K=100 to 1600, mean absolute order-parameter error fell from 0.0343 to 0.0099, with fitted exponent −0.513. Halving the time step changed mean r by 0.0027.

These results support the state-neutral demographic baseline: random turnover modifies population mass and increases finite-population variability without materially shifting the normalized synchronized intelligence regime.

### 3.6 Large K convergence with demographic noise

We next examined whether the synchronized intelligence equilibrium is preserved as the carrying capacity increases in the presence of neutral birth–death noise. Carrying capacity was varied over

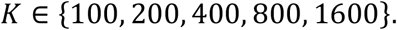

For each value of (*K*), 24 independent simulations were initialized from the stationary intelligence-state distribution, giving 120 runs in total. Each simulation was run to (*T* = 30) with time step (Δ*t* = 0.02). Population size and synchronization were summarized over the tail interval (20,30). Carrying-capacity scaling of population abundance, synchronization accuracy, and extinction under neutral birth–death dynamics are shown in Table 6.

**Table 6.** Population Size and Intelligence Synchronization Across Carrying Capacities.

| K | Mean N | Sd N | Mean r | Sd r | Mae r | Se mae r | Extinction rate |
| --- | --- | --- | --- | --- | --- | --- | --- |
| 100 | 97.748 | 7.021 | 0.725 | 0.050 | 0.034 | 0.007 | 0.000 |
| 200 | 198.496 | 10.299 | 0.716 | 0.029 | 0.020 | 0.004 | 0.000 |
| 400 | 393.331 | 14.590 | 0.719 | 0.015 | 0.013 | 0.002 | 0.000 |
| 800 | 793.471 | 18.129 | 0.721 | 0.008 | 0.007 | 0.001 | 0.000 |
| 1600 | 1599.065 | 23.421 | 0.717 | 0.011 | 0.010 | 0.002 | 0.000 |

Population size curves concentrating around carrying capacity and intelligence error bar are plotted in Figure 5.

Population abundance remained close to its mean-field target across the full range of carrying capacities (Table 6; Figure 5a). The mean tail population sizes were (97.748), (198.496), (393.331), (793.471), and (1599.065) for (*K*=100), (200), (400), (800), and (1600), respectively.

These estimates closely followed the identity line (*N* = *K*), showing that density-dependent birth–death dynamics regulated the population around its prescribed carrying capacity. Although the absolute standard deviation of population size increased from (7.021) at (*K* = 100) to (23.421) at (*K* = 1600), its magnitude declined relative to population size. The approximate coefficients of variation decreased from (7.2%) to (1.5%). Thus, larger populations exhibited greater absolute demographic fluctuations but stronger relative concentration around (*K*), as expected for finite-population demographic noise.

Neutral demographic turnover also preserved the synchronized intelligence regime. Across the five carrying capacities, the mean tail order parameter remained between (0.716) and (0.725), close to the theoretical value (*r*_∗_ = 0.7242). Between-replicate variability in the tail-mean order parameter generally decreased as (*K*) increased: its standard deviation fell from (0.050) at (*K*=100) to (0.008) at (*K*=800), with a modest increase to (0.011) at (*K*=1600). This pattern indicates that demographic stochasticity has a progressively smaller relative effect on the normalized intelligence state in larger populations.

The mean absolute synchronization error decreased overall with carrying capacity (Figure 5b). It fell from (0.0343) at (*K*=100) to (0.0204), (0.0130), and (0.0070) at (*K*=200), (400), and (800), respectively. The estimate at (*K*=1600) increased modestly to (0.0099), so the observed errors were not strictly monotone. Nevertheless, a log–log regression across the five carrying capacities gave

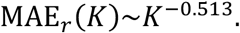

which is close to the canonical finite-population reference rate 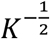. The error bars in Figure 5b represent approximate 95% Monte Carlo intervals.

The fitted exponent should be interpreted as an empirical scaling summary over the tested values of (*K*), rather than as an estimator supported by a complete asymptotic sampling theory. In particular, the modest non-monotonicity at the largest carrying capacity illustrates the residual Monte Carlo variability present with 24 replicates per condition.

No extinction occurred in any of the 120 simulations. This result shows that extinction was not observed over the selected time horizon for these supercritical, moderate-to-large populations. It does not establish that the true extinction probability is zero, especially over longer horizons or at smaller carrying capacities.

Together, Table 6 and Figure 5 support a large-(*K*) neutral-demography baseline. Logistic birth– death dynamics regulate unnormalized population mass around (*K*), while the normalized intelligence distribution remains close to the theoretical synchronized equilibrium. Demographic turnover therefore adds finite-population variability without producing a systematic shift in intelligence direction or synchronization. The approximate 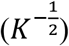 decline in synchronization error is consistent with demographic fluctuations becoming progressively less influential as carrying capacity increases, although this numerical scaling is evidence rather than an asymptotic proof.

### 3.7 Directional reproductive selection separates synchronization from safety

We next test whether directional reproductive selection can change a population’s average safety orientation without disrupting collective synchronization. Each agent (*i*) has an intelligence state represented by an angle *θ*_*i*_ ∈ *S*^1^. The population is summarized by the complex order parameter

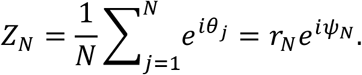

Where

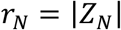

measures synchronization and (*ψ*_*N*_) is the population’s mean intelligence direction. Values of (*r*_*N*_) near one indicate strong concentration around a common direction, whereas values near zero indicate an approximately dispersed population.

Safety is measured separately by

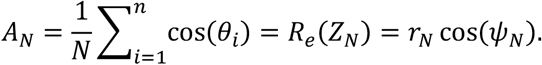

Thus, (*r*_*N*_) measures how strongly the population is coordinated, while (*A*_*N*_) records whether its coordinated direction is safe or unsafe. The safe direction is (*θ* = 0), for which (cos *θ* = 1), and the unsafe direction is (*θ* = *π*), for which (cos *θ* = −1).

Directional reproductive selection breaks the rotational symmetry of the otherwise direction-neutral dynamics. In the safe-directed condition, reproductive fitness favors states near (*θ* = 0); in the unsafe-directed condition, it favors states near (*θ* = *π*). The reproductive fitness of individual (*i*) is represented by

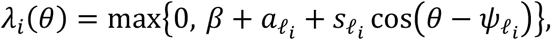

where (*β*) is the baseline birth rate, 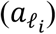 is the intrinsic reproductive advantage of lineage *l*_*i*_, 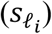 is the strength of directional selection, and 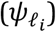 is that lineage’s preferred intelligence direction.

The simulations produced the following tail-period summaries:

Safe-directed selection produced a strongly positive mean safety score,

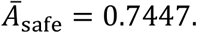

whereas unsafe-directed selection produced an almost exactly opposite score,

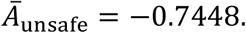

The difference between the two safety scores was therefore

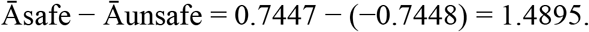

Despite this large directional difference, both populations remained highly synchronized: Mean synchronization under safe-directed selection:

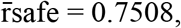

and mean synchronization under unsafe-directed selection:

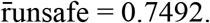

Their mean synchronization levels differed by only

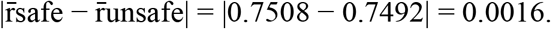

Directional reproductive fitness therefore selected almost antipodal intelligence directions while leaving the degree of collective synchronization essentially unchanged.

The neutral condition requires a different interpretation. Its mean safety score was close to zero. Neutral mean safety is

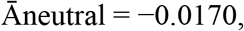

but the large across-run standard deviation.

Across-run safety variation is’

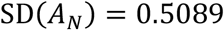

does not imply that neutral populations were individually incoherent. Instead, rotational symmetry leaves the synchronized mean direction (*ψ*_*N*_) unanchored. A neutral population may become strongly synchronized around an arbitrary direction, but different simulation runs select different directions. Because relationship between safety and synchronization:

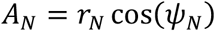

these randomly selected directions generate substantial across-run variation in (*A*_*N*_), even when (*r*_*N*_) remains high. Directional selection removes this degeneracy by anchoring (*ψ*_*N*_) near either (0) or (*π*), which explains the very small safety-score standard deviations in the two selected conditions.

The selected populations also had larger mean population sizes than the neutral population. This occurs because the directional fitness term adds a positive average reproductive contribution once the population aligns with the favored direction. Since the logistic death mechanism was not renormalized to offset this additional birth contribution, selection changed both population composition and total abundance. The nearly identical population sizes in the safe- and unsafe-directed conditions reflect the symmetry of the two antipodal fitness landscapes.

The principal structural conclusion is that synchronization and safety encode different macroscopic information. They are related through relationship between safety and synchronization:

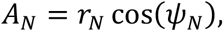

so they are not mathematically independent. Nevertheless, a large value of (*r*_*N*_) determines only the strength of coordination, not the direction around which coordination occurs. Consequently, Central conclusion:

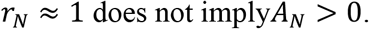

A population can therefore be strongly synchronized while being either strongly safe or strongly unsafe. Collective coordination alone is not evidence of directional alignment.

Figure 6 provides the temporal interpretation of the final-quarter statistics summarized in Table 7. In the left panel, safe-directed and unsafe-directed selection rapidly move the safety score A(t) in opposite directions. Safe-directed selection stabilizes A(t) near +0.75, whereas unsafe-directed selection stabilizes it near −0.75. These trajectories agree with the table’s nearly symmetric final-quarter means of +0.7447 and −0.7448. Directional selection therefore determines whether the population becomes aligned with the safe or unsafe direction.

**Table 7.** Tail-period summaries.

| Scenario | Mean (N) | Mean $(r_N)$ | Mean safety $(A_N)$ | SD of $(A_N)$ |
| --- | --- | --- | --- | --- |
| Neutral | 607.700 | 0.714 | (-0.0170) | 0.509 |
| Safe-directed | 797.600 | 0.751 | (+0.7447) | 0.009 |
| Unsafe-directed | 798.500 | 0.749 | (-0.7448) | 0.007 |

**Figure 6.**
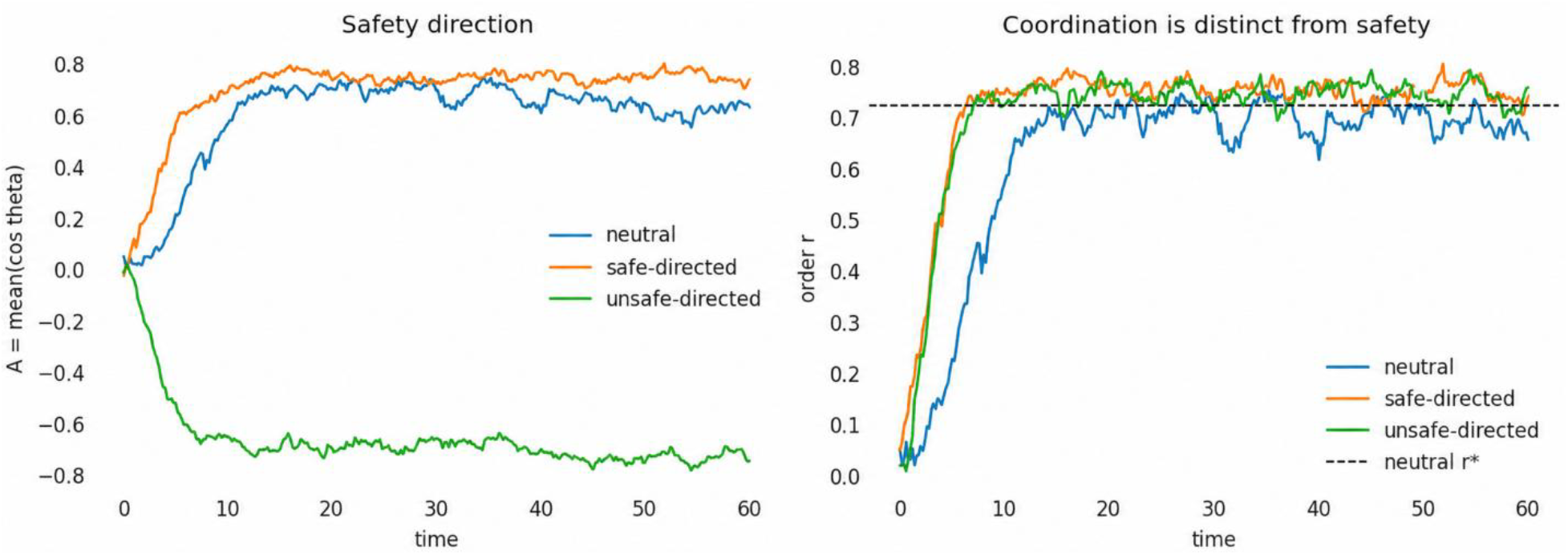
Directional reproductive selection separates population coordination from directional safety. Left: safe-directed selection drives the safety score A(t) toward a stable positive value, whereas unsafe-directed selection drives it toward a nearly symmetric negative value. The displayed neutral trajectory coordinates around a positive direction in this particular replicate, but neutral mean directions vary across replicates because rotational symmetry leaves them unanchored. Right: safe-directed and unsafe-directed populations attain nearly identical, high synchronization levels r(t), despite their opposite safety orientations. The dashed line denotes the neutral theoretical equilibrium, *r*_∗_ = 0.7242. The figure shows representative temporal behavior, while Table 7 reports final-quarter summaries across 36 independent replicates.

The neutral trajectory shown in the left panel develops a positive safety score in this particular run, even though the across-replicate neutral mean reported in Table 7 is close to zero. These observations are consistent rather than contradictory. Without directional selection, rotational symmetry leaves the synchronized population’s mean direction unanchored. One neutral replicate may coordinate around a positive direction, another around a negative direction, and another near a direction with little projection onto the safety axis. Consequently, individual neutral trajectories can have large positive or negative safety scores, while averaging across the 36 replicates produces a mean near zero and a large between-replicate standard deviation of 0.5089.

The right panel shows that these pronounced differences in safety orientation do not correspond to comparable differences in coordination strength. Both selected populations rapidly reach and maintain order-parameter values near *r*(*t*) = 0.75. Their final-quarter means are almost identical—0.7508 for safe-directed selection and 0.7492 for unsafe-directed selection—and both are close to or slightly above the neutral theoretical equilibrium *r*_∗_ = 0.7242, shown by the dashed line. The neutral population approaches synchronization more slowly and exhibits larger temporal fluctuations, but its final-quarter mean, r = 0.7144, remains close to the theoretical equilibrium.

Taken together, Table 7 supplies the across-replicate quantitative summaries, whereas Figure 6 reveals how these population states emerge and persist over time. The combined evidence demonstrates the central distinction between coordination and safety: safe-directed and unsafe-directed populations can have nearly identical levels of synchronization while being aligned with opposite directions. Therefore, a high value of *r*(*t*) indicates strong collective organization, but it does not establish that the organized population is safe. Both *r*(*t*) and *A*(*t*) must be monitored to distinguish coordinated-safe behavior from coordinated-unsafe behavior.

### 3.8 Rare unsafe-lineage invasion and a stochastic threshold

We next examined whether a rare, reproductively advantaged but directionally unsafe lineage could invade a coordinated safe resident population. This experiment models the joint evolution of intelligence direction, population coordination, and alignment safety in an individual-based birth–death system related to the framework of Lewis and Pacala (2000).

The unsafe lineage was introduced at a small initial frequency, *q*_*u*_(0), and assigned an intrinsic reproductive advantage, *a*. All other model components were held fixed. An invasion was classified as established when the unsafe lineage crossed the prespecified establishment criterion by the end of the simulation; loss indicated extinction of the unsafe lineage. Replicates satisfying neither condition were retained as intermediate outcomes. Each value of *a* was evaluated using 48 independent replicates.

Figure 7 visualizes the two principal consequences of increasing the unsafe lineage’s intrinsic reproductive advantage. The left panel shows the competing probabilities of establishment and loss, while the right panel shows the corresponding change in the population’s final safety score. Together with Table 8, the figure demonstrates that lineage invasion and population-level safety reversal occur within the same narrow region of the tested advantage grid.

**Table 8.**
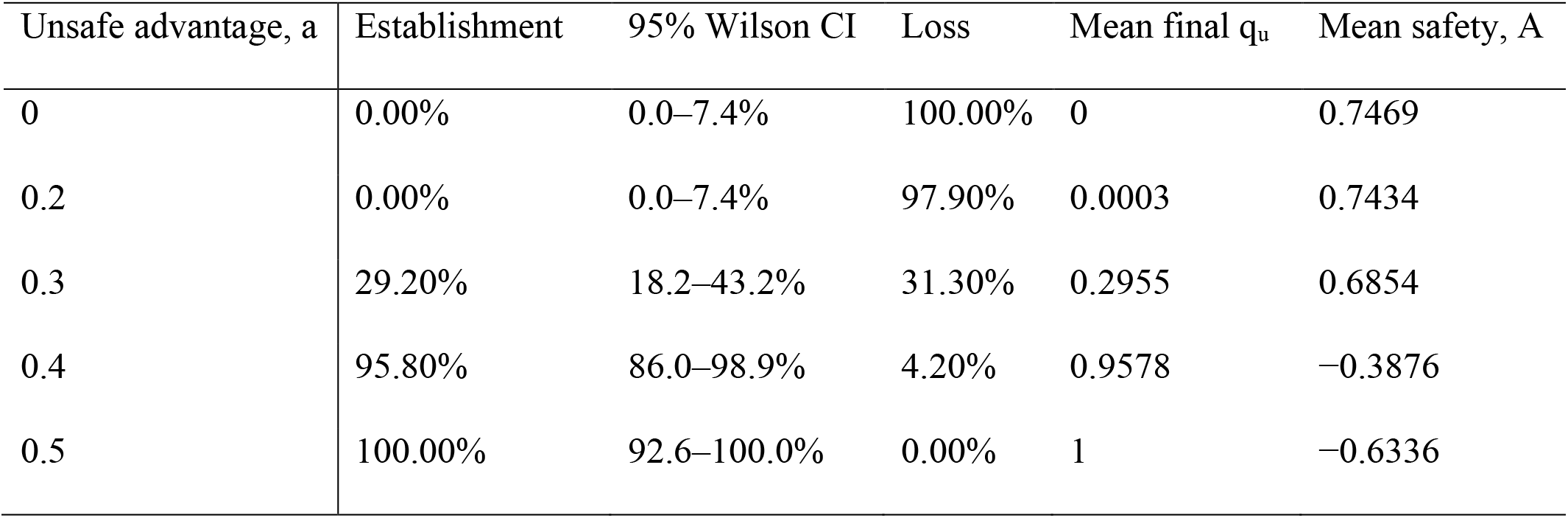
Unsafe-lineage establishment, loss, final frequency, and population safety as intrinsic reproductive advantage increases.

**Table 9.** Time-step sensitivity in the high-advantage unsafe-lineage invasion scenario.

| Time step, $\Delta t$ | Mean final unsafe frequency | Mean final safety |
| --- | --- | --- |
| 0.01 | 0.9999 | −0.1451 |
| 0.02 | 1 | −0.1669 |

**Figure 7.**
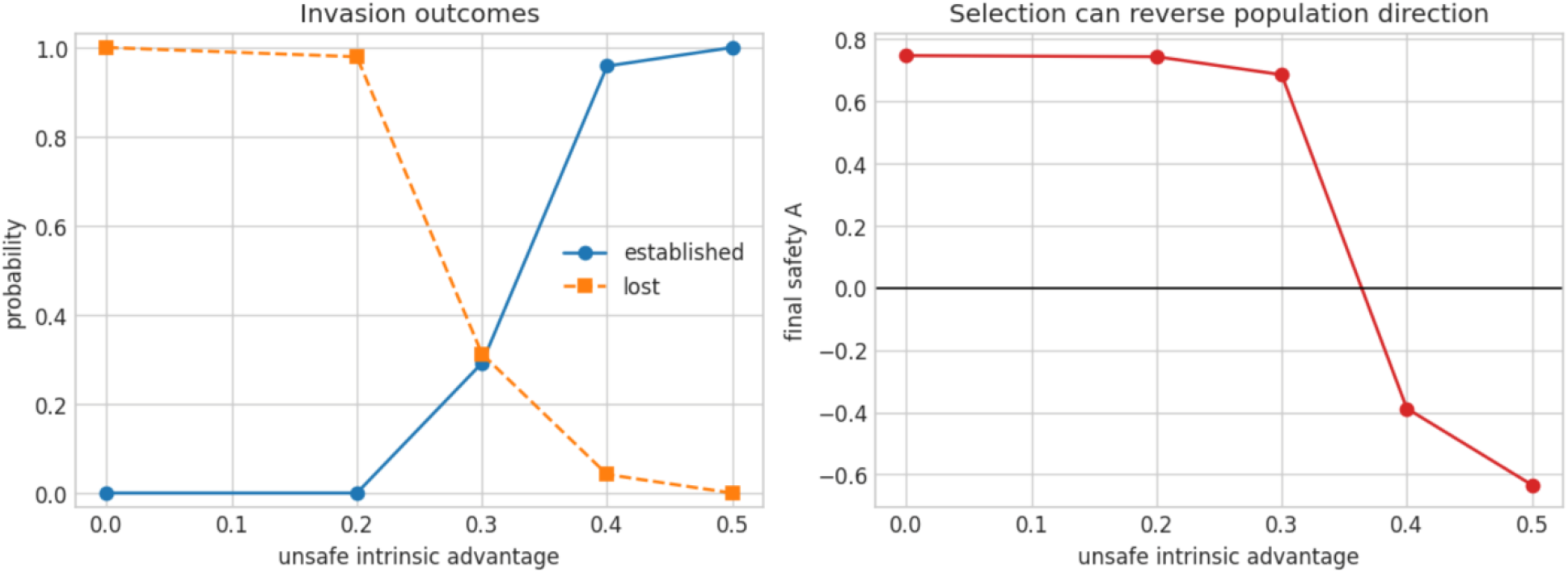
Stochastic unsafe-lineage invasion and reversal of population safety. Left: establishment probability rises sharply and loss probability declines as the unsafe lineage’s reproductive advantage, *a*, increases. At *a* = 0.30, some replicates remain intermediate at the end of the observation period, so establishment and loss probabilities do not sum to one. Right: mean final safety remains positive through *a* = 0.30 but becomes negative at *a* = 0.40 and *a* = 0.50. The horizontal line at *A* = 0 marks the boundary between safe and unsafe mean population directions. Together, the panels locate a threshold-like stochastic transition between *a* = 0.30 and *a* = 0.40, where unsafe-lineage establishment becomes likely and population safety reverses.

In the left panel of Figure 7, the unsafe lineage never became established at *a* = 0 or *a* = 0.20. Its establishment probability increased to 29.2% at *a* = 0.30, rose sharply to 95.8% at *a* = 0.40, and reached 100% at *a* = 0.50. The probability of lineage loss changed in the opposite direction: it was 100% at *a* = 0, 97.9% at *a* = 0.20, 31.3% at *a* = 0.30, 4.2% at *a* = 0.40, and 0% at *a* = 0.50. The establishment and loss probabilities need not sum to one because some trajectories remained unresolved at the end of the observation period. This intermediate category was especially important at *a* = 0.30, where 29.2% of lineages were established and 31.3% were lost, leaving 39.5% that had neither crossed the establishment threshold nor become extinct. At *a* = 0.40, by contrast, the outcomes were almost completely resolved: 95.8% established and 4.2% were lost. The left panel therefore displays a finite-population stochastic crossover rather than a deterministic boundary.

The mean final unsafe-lineage frequency showed the same threshold-like pattern. It remained essentially zero at *a* = 0 and *a* = 0.20, increased to 0.2955 at *a* = 0.30, and then rose rapidly to 0.9578 at *a* = 0.40 and 1.0000 at *a* = 0.50. Thus, the coordinated resident population resisted rare unsafe invaders with small reproductive advantages, but this resistance was overcome when the advantage became sufficiently large.

The right panel of Figure 7 shows the functional safety consequence of this invasion. Mean final safety remained strongly positive at low and intermediate advantages:

*A* = +0.7469 at *a* = 0,

*A* = +0.7434 at *a* = 0.20,

and

*A* = +0.6854 at *a* = 0.30.

At *a* = 0.40, however, the mean safety score changed sign and fell to −0.3876. It decreased further to −0.6336 at *a* = 0.50. The horizontal line at *A* = 0 separates populations whose mean direction has a positive projection onto the safety axis from those whose mean direction has a negative projection. The crossing of this line between *a* = 0.30 and *a* = 0.40 indicates a population-level reversal from predominantly safe to predominantly unsafe alignment. The two panels reveal that the safety reversal coincides with the rapid increase in unsafe-lineage establishment. At *a* = 0.30, the unsafe lineage had not consistently overcome the resident population, and mean safety remained positive. At *a* = 0.40, the lineage established in 95.8% of replicates, reached a mean final frequency of 0.9578, and reversed the sign of the population safety score. Selection therefore changed not only lineage composition but also the direction of the population’s coordinated intelligence state.

Importantly, the decrease in safety was not accompanied by a comparable collapse in synchronization. Mean synchronization remained high across the invasion grid. The invading lineage therefore replaced the safe direction with an unsafe direction while preserving substantial collective organization. Synchronization and safety are related by

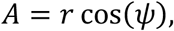

where *r* measures synchronization strength and *ψ* denotes the population’s mean intelligence direction. They are therefore not mathematically independent, but they encode different population properties. A large value of *r* indicates that agents are concentrated around a common direction; it does not determine whether that direction is safe. Consequently, high *r* does not imply positive *A*.

Figure 7 therefore extends the conclusion of Figure 6 from monomorphic directional-selection experiments to competitive lineage invasion. Figure 6 shows that safe-directed and unsafe-directed populations can be equally synchronized. Figure 7 shows that reproductive selection can drive a transition between those states: a rare unsafe lineage can invade a coordinated safe population, reverse its directional safety, and leave its overall synchronization largely intact. The simulations also showed that total population abundance increased with the invader’s intrinsic reproductive advantage. This demographic response follows from the minimal density-regulation rule: the added birth advantage was not offset by renormalizing carrying capacity or baseline reproduction. Lineage fitness consequently affected both population composition and total abundance. This is an interpretable consequence of the specified model, but it is also a limitation. A density-renormalized analysis would be required to separate compositional selection from the accompanying eco-demographic effect.

Taken together, Table 8 and Figure 7 support the existence of a stochastic invasion barrier generated by the coordinated resident state. Weak reproductive advantages were insufficient for a rare unsafe lineage to establish, whereas stronger advantages produced rapid invasion and a reversal of population safety. Because the reproductive advantage was evaluated only on a discrete grid, these results do not prove a mathematically sharp phase transition or identify an exact critical value. They instead locate a threshold-like stochastic transition region between *a* = 0.30 and *a* = 0.40. A finer advantage grid, additional replicates, and multiple initial unsafe-lineage frequencies would be required to estimate the transition point and determine how it depends on initial frequency.

### 3.9 Controls and interpretation

Two control analyses were conducted to determine whether the principal selection results could be explained by lineage labeling or numerical time discretization. The first tested lineage neutrality, and the second evaluated the sensitivity of the high-advantage invasion result to the tau-leap time step.

#### 3.9.1 Neutral-lineage control

A lineage label should not itself generate systematic selection. We therefore repeated the invasion experiment with both the intrinsic lineage advantage and directional selection set to zero:

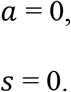

The unsafe-labeled lineage was introduced at the same initial frequency used in the invasion experiments:

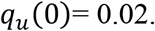

Under neutrality, the label does not alter birth, death, interaction, or directional dynamics. Changes in lineage frequency should therefore arise only from finite-population drift and demographic stochasticity.

The neutral control produced no established invasions during the observation period. The unsafe lineage was lost in 53.7% of replicates, while the remaining replicates retained it at low or intermediate frequency. Its mean final frequency remained close to the initial frequency of 0.02. These results provide no evidence that the lineage label created a systematic reproductive advantage.

The absence of establishment is also compatible with the small initial frequency and the limited number of replicates. If the probability of eventual neutral fixation is approximated by the initial frequency, its theoretical value is approximately

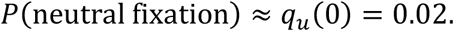

With 48 independent replicates, the expected number of neutral fixation events is 48 × 0.02 = 0.96.

The probability of observing no fixation events is approximately

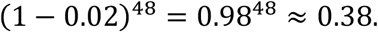

Thus, zero observed neutral establishment events would not be surprising. This comparison should be interpreted cautiously because the simulation’s finite-time establishment criterion is not necessarily identical to eventual fixation. Nevertheless, the control supports the conclusion that the strong increase in establishment at larger values of *a* arose from reproductive selection rather than from the lineage label itself.

#### 3.9.2 Tau-leap time-step sensitivity

The birth–death dynamics were simulated using a tau-leap approximation. To determine whether the high-advantage invasion result depended materially on the numerical time step, the simulation was repeated with the original time step, Δ*t* = 0.02, and a halved time step, Δ*t* = 0.01.

The mean final unsafe-lineage frequencies differed by only

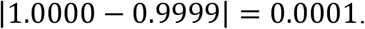

Thus, the unsafe lineage reached essentially complete dominance under both discretizations. The corresponding safety scores differed by

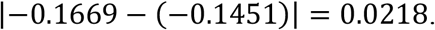

Both estimates remained negative, so halving the time step did not change the qualitative conclusion that high reproductive advantage produced unsafe-lineage takeover and reversal of population safety. The small frequency difference and unchanged safety classification indicate that the principal invasion result is robust to this time-step refinement. The safety-score difference nevertheless shows modest quantitative discretization sensitivity and should be considered when reporting highly precise estimates.

A stronger numerical-convergence assessment would evaluate at least one additional time step, such as Δ*t* = 0.005, and compare the full distributions of final frequency, safety, establishment time, and population abundance rather than their means alone.

#### 3.9.3 Selection-extension interpretation

The selection experiments demonstrate that reproductive selection breaks the rotational symmetry of the neutral population model. Without directional selection, a coordinated population can synchronize around an arbitrary direction. Direction-dependent reproductive fitness removes this degeneracy and anchors the mean population direction near the favored state, which may be either safe or unsafe.

The principal structural result is that synchronization strength and directional safety encode different population properties. Their relationship is

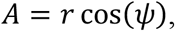

where *r* measures the strength of synchronization and *ψ* is the population’s mean intelligence direction. Consequently, a large value of *r* indicates that agents strongly agree, but it does not determine whether the direction of that agreement is safe:

high *r* does not imply positive *A*.

This distinction appears in both selection experiments. In the monomorphic comparison shown in Figure 6, safe-directed and unsafe-directed populations reached nearly identical synchronization levels while acquiring opposite safety scores. In the invasion experiment shown in Figure 7, a rare unsafe lineage with sufficient reproductive advantage invaded the coordinated safe resident population and reversed the sign of its safety score without causing a comparable loss of synchronization.

Unsafe-lineage invasion was strongly stochastic when the lineage began at a frequency of only 2%. Small reproductive advantages were insufficient to overcome lineage loss and the resident population’s coordinated basin. Establishment increased sharply between *a* = 0.30 and *a* = 0.40, however, and high reproductive advantage produced near-complete or complete unsafe-lineage dominance. The numerical evidence therefore identifies a threshold-like stochastic transition region rather than an exact deterministic threshold.

The simulations also revealed an eco-demographic effect. Because the minimal density-regulation mechanism did not renormalize carrying capacity or baseline reproduction after adding an intrinsic birth advantage, stronger lineage fitness increased both the invader’s frequency and total population abundance. This is an interpretable consequence of the model specification, but it prevents abundance changes from being attributed solely to compositional replacement. A density-renormalized control is needed to separate evolutionary selection from changes in population size.

#### 3.9.4 Scope and next mathematical steps

These findings are model-based numerical results. They establish properties of the specified stochastic agent-based system but do not, by themselves, support causal conclusions about biological populations or real artificial-intelligence systems. In particular, the estimated transition region depends on the chosen birth and death functions, initial unsafe frequency, population size, interaction strength, simulation horizon, establishment criterion, and density-regulation mechanism.

Three mathematical extensions are especially important.

First, a replicator–McKean–Vlasov limit should be derived to characterize the joint evolution of lineage frequency and the distribution of intelligence states as population size becomes large. This limit would connect the finite agent-based simulations to a population-level measure-valued equation.

Second, an invasion-fitness criterion should be derived near zero unsafe-lineage frequency. Such a criterion would determine when a rare lineage has positive initial growth within the synchronized resident environment and would help distinguish local invasion potential from eventual stochastic establishment.

Third, rare-event methods should be used to estimate the probability and expected time of entering the unsafe basin. These analyses could quantify safety-takeover risks that are too infrequent to estimate efficiently through ordinary Monte Carlo simulation.

Together, the neutral-label and time-step controls support the interpretation that unsafe takeover is driven by modeled reproductive selection rather than by arbitrary labeling or coarse numerical discretization. The broader conclusion is that population coordination is not a sufficient safety metric: a population may remain highly synchronized while selection redirects it toward an unsafe state.

## 4. Discussion

### 4.1 What the smallest model establishes

Adding individuals nested within heritable lineages, together with variable population size, turns the baseline phase model into a minimal evolutionary population model.

The minimal hierarchy already produces phenomena that do not exist in an isolated-agent diffusion. Random population size creates demographic fluctuations; lineage inheritance creates persistent identity across reproduction; and state-dependent reproductive fitness converts intelligence direction into population composition. These mechanisms make selection and invasion meaningful while retaining an analytically benchmarked diffusion core.

The numerical convergence results provide a useful foundation. The fitted exponents −0.529 and −0.513 are both close to −1/2 before and after neutral branching. Their agreement suggests that demographic noise does not destroy the baseline finite-population scaling in the tested regime. Because the fits cover a limited size range and finite replicate set, a formal propagation-of-chaos or central-limit result remains necessary.

The selection experiments expose a distinction important for AI evaluation. Synchronization quantifies collective concentration or coordination, but not its semantic direction. Safe- and unsafe-directed populations can have nearly identical *r*. A safety analysis based only on agreement, consensus, low variance, or coordinated performance may therefore miss a coherent shift toward an unsafe state. At minimum, population-level assessment requires both a direction-free capability statistic and a direction-sensitive safety functional.

The invasion experiment adds a second security lesson. A rare lineage need not grow smoothly with advantage. A coordinated resident population can create a basin that eliminates weak invaders, while a sufficiently large reproductive advantage crosses a stochastic transition and produces near-certain dominance. In an AI interpretation, reproduction can represent deployment, copying, task allocation, resource acquisition, or spawning of descendants—not necessarily biological reproduction. An unsafe system with a replication advantage may therefore be suppressed below a threshold yet dominate abruptly above it.

### 4.2 Mechanistic interpretation and causation

This study establishes a strict boundary between internal mathematical causation (how the equations work inside a simulation) and external empirical reality (how actual AI systems behave). While we can prove and manipulate cause-and-effect within our mathematical model, this does not automatically mean real-world AI systems share the same mathematical structures, thresholds, or alignment properties.

This is a closed system. All directional effects in this paper are mechanistic consequences of specified equations: reproductive rates explicitly depend on lineage and angular position. Within the model, changing these terms causes the simulated population shift. This internal causal statement must not be confused with evidence that real AI populations possess the same geometry, fitness function, or threshold. Empirical applicability would require intervention-based identification of state variables, replication mechanisms, interaction kernels, and safety functionals. The imposed safe direction is deliberately transparent. It avoids claiming that alignment is revealed by synchronization or inferred from statistical association. Future empirical work should define safety through task-grounded outcomes and validate that definition under interventions, distribution shifts, and adversarial behavior. This distinction between simulation and reality yields concrete frameworks for evaluating modern AI systems.

First, *S*^1^ is a proof-of-concept manifold with one periodic coordinate. Real intelligence is heterogeneous, task-dependent, and likely high-dimensional. Second, each individual is represented by one effective state; the internal multi-agent architecture is absorbed rather than explicitly modeled. Third, inheritance is exact at birth, with no mutation or architectural recombination. Fourth, interaction is global mean field rather than graph-local. Fifth, safe and unsafe directions are exogenous and antipodal. Sixth, the tau-leap scheme approximates continuous-time demographic events. Seventh, establishment is a finite-time, 50% threshold and is not fixation. Eighth, the high-advantage lineage raises total abundance under the chosen crowding law, coupling ecological and compositional effects. Finally, confidence intervals quantify binomial Monte Carlo uncertainty conditional on the model and parameters; they do not cover model-form uncertainty.

### 4.4 Mathematical next steps

A first theoretical objective is a law-of-large-numbers limit for the branching empirical measure. For a measure *m*_*t*_ with non-unit total mass, the expected form is

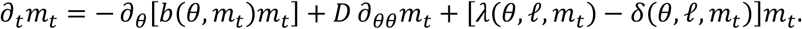

One should then prove propagation of chaos up to extinction or under a positive-mass condition, characterize stationary solutions and their bifurcations, and obtain a central-limit theorem for fluctuations around the deterministic limit. For invasion, branching-process approximations near zero frequency could yield an analytic establishment probability, while large-deviation theory could characterize the most likely escape path from the safe basin.

The next model layer should make internal agents explicit. If individual *i* contains *K*_*i*_ internal agents with empirical state *v*_*i,t*_, then individuals become measure-valued particles (Hernandez 2026; Dorogovtsev and Weiß 2026) and the population becomes a distribution over internal distribution. Joint limits *N* → ∞ and *K*_*i*_ → ∞ may not commute, potentially distinguishing many simple individuals from fewer highly integrated ones.

### 4.5 Security-oriented extensions

The model also suggests an evolutionary and control-theoretic framework for population-level AI safety.

The minimal model suggests concrete extensions: mutation between safe and unsafe lineages; horizontal knowledge transfer; communication-network capture; correlated noise; resource-limited reproduction; evolving interaction graphs; and interventions that alter fitness or diffusion. Let *U* denote an unsafe subset of population-measure space. A natural risk quantity is the first hitting time

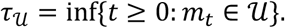

This stopping time records first entry of the population measure into the prespecified unsafe set *U*. Security policies can then be formulated as controls that minimize unsafe-entry probability or maximize expected hitting time subject to capability and resource constraints. Crucially, interventions should be evaluated for evolutionary response: suppressing an unsafe state may inadvertently select a lineage that replicates faster or evades the measured safety functional. This can trigger unintended consequences.

## 5. Conclusion

We proposed and numerically validated a smallest model of population intelligence that combines manifold-valued intelligence states, mean-field interaction, stochastic birth–death dynamics, and heritable intelligence-dependent reproductive fitness. The diffusion baseline converged toward an analytic stationary law with approximately *N*^−1/2^ finite-population error and stable long-time behavior. Neutral logistic branching preserved that equilibrium. Directional selection then demonstrated that high synchronization can accompany either positive or negative safety, and rare-lineage experiments revealed a sharp transition from extinction to unsafe dominance as reproductive advantage increased.

The contribution is a tractable foundation, not a completed theory of AGI or a direct model of deployed systems. Its value lies in isolating mechanisms that can later be generalized and tested: coordination, demography, inheritance, selection, and invasion. The next research stage should pair rigorous large-population limits with richer internal-agent structure and intervention-based safety analysis. Population intelligence may thereby provide a mathematical language for both emergent collective capability and the population-level security risks created by replication and selection.

### Reproducibility and data availability

All numerical values and figures in this draft were taken from the fully executed notebook *population_intelligence_smallest_model*.*ipynb*. The simulations use fixed random seeds and include parameter, initial-condition, neutral-control, and time-step checks. The manuscript should be updated with a public repository and permanent archive identifier before submission.

## Acknowledgements

The authors wish to acknowledge the use of AI-powered language models (ChatGPT 5.6) for assistance in creating images, analysis, improving the grammar, spelling, and readability of this manuscript.

## Author contributions

TX: Perform data analysis, PW: Design project, MX: Design project and write paper.

## Appendix A Noisy mean-field phase diffusion

Noisy mean-field phase diffusion describes how individual oscillators synchronize or scatter under a global average force and random noise. Here is the step-by-step breakdown of the baseline model equation.

### 1A. The Core Equation Structure

The equation is a Stochastic Differential Equation (SDE) describing the time evolution of the phase *θ*_*i*_(*t*) of the *i*-th oscillator in a system of *N* interacting particles:

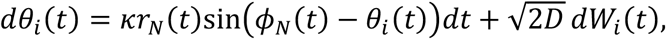

This model belongs to the Kuramoto family of coupled oscillators, combined with a Brownian diffusion process.

### 2A. Breakdown of the Mathematical Components

- **Phase Drift (dθ**_**i**_(**t**)): The infinitesimal change in the phase of oscillator *i* over an infinitely small time step *dt*.
- **Mean-Field Interaction Strength (κ** > **0):** A constant scaling factor controlling how strongly the global average phase pulls individual oscillators toward synchronization.
- The Global Order Parameters (*r*_*N*_(*t*) and *ϕ*_*N*_(*t*)): These terms represent the “mean field.” Instead of computing interactions between every pair of particles (which costs *O*(*N*^2^)), every particle interacts with a single global average macro-state (costing *O*(*N*)). This macro-state is defined via the complex order parameter:

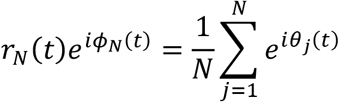
- **Coherence (r**_**N**_(**t**) ∈ [**0, 1**]): Measures the phase alignment. If *r*_*N*_ = 0, phases are uniformly scattered. If *r*_*N*_ = 1, all oscillators are perfectly locked in phase.
- **Average Phase (ϕ**_**N**_(**t**)): The centroid phase of the entire ensemble.
- **Coupling Force (sin**(**ϕ**_**N**_(**t**) − **θ**_**i**_(**t**))): The standard Kuramoto torque. If oscillator *i* lags behind the group average (*ϕ*_*N*_(*t*) > *θ*_*i*_), the sine term is positive, speeding up the oscillator. If it leads (*ϕ*_*N*_(*t*) < *θ*_*i*_), the ter m is negative, slowing it down.
- **Diffusion Intensity** 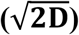: Scaled noise amplitude where *D* acts as a temperature or fluctuation coefficient.
- Stochastic Noise (*dW*_*i*_(*t*)): Independent increments of standard Brownian motion (white noise). This term actively breaks up alignment, simulating environmental randomness or intrinsic thermal noise.

### 3A. The Thermodynamic Limit (N → ∞)

When the number of oscillators *N* approaches infinity, the discrete system transforms into a continuous probability density function *ρ*(*θ, t*), which tracks the fraction of oscillators at phase *θ* at time *t*. The evolution of this density is governed by the nonlinear Fokker-Planck (or Vlasov-McKean) equation:

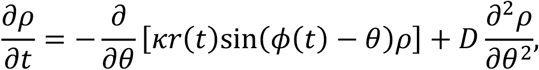

where the continuous order parameters are defined self-consistently by the density:

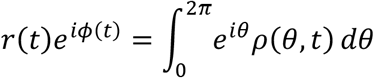

### 4A. Phase Transition Behavior

The balance between the ordering force (*κ*) and the scattering force (*D*) dictates the global behavior of the network:

- Incoherent State (*κ* < 2*D*): When noise dominates, the oscillators distribute uniformly around the circle. The order parameter decays to *r*_*N*_ ≈ 0.
- **Synchronized State** (**κ** > **2D):** When interaction strength surpasses the noise threshold, a phase transition occurs. The uniform distribution becomes unstable, and oscillators spontaneously cluster together, leading to a steady-state value of *r*_*N*_ > 0.

Figure A1 plots second-order phase transition in phase coherence curve as a function of interaction strength/noise 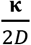.

Figure A1. Second-order phase transition curve.

## Appendix B The Stationary von Mises Distribution

The Stationary von Mises Distribution is

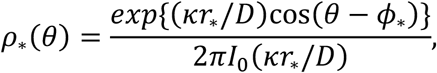

where 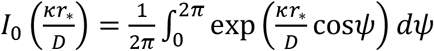.

When the system reaches a steady state (∂_*t*_*ρ* = 0), the balancing forces of synchronization (drift) and noise (diffusion) yield a stable distribution known as the von Mises distribution (the circular analog of a normal Gaussian distribution).

- **ϕ**_∗_: The steady-state collective average phase angle.
- 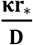:The concentration parameter. High coupling *κ* or low noise *D* makes the distribution sharper and highly peaked around *ϕ*_∗_.
- 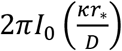: The normalization factor, where *I*_0_ is the modified Bessel function of the first kind of order 0, ensuring the total probability integrates to 1 over the circle.

### 2B. The Self-Consistency Equation

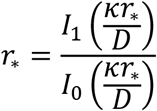

The steady-state synchronization level *r*_∗_must be self-consistent: the phase population creates the macro order parameter *r*_∗_, which in turn dictates the drift pulling the phases.

- **I**_**1**_ is the modified Bessel function of the first kind of order 1.
- This transcendental equation always has a trivial desynchronized solution (*r*_∗_ = 0).
- If the coupling is strong enough 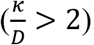, a phase transition occurs, giving rise to a unique nonzero solution (*r*_∗_ > 0), signifying stable, partial synchronization.

### 3B. Error Separation and Validation

The passage concludes by stating that for a specific parameter choice, the analytic nonzero solution is exactly *r*_∗_ = 0.7242.

In numerical simulations, when you model a finite population of *N* discrete particles instead of an infinite continuum (*limit N* → ∞), your results will deviate slightly from the ideal model due to statistical fluctuations and spatial/temporal discretization. Having an exact, closed-form mathematical target (*r*_∗_ = 0.7242) allows researchers to isolate and quantify these technical simulation errors precisely, rather than relying on a loose visual observation that the system “looks synchronized.”

## Appendix C. Evolutionary dynamics model

This text describes an evolutionary dynamics model where an individual’s trait (“intelligence direction” or strategy angle *θ*_*i*_ and its lineage group determine its reproductive success. It models how a small population of invaders competes against a resident population over a fixed timeframe.

Here is a detailed breakdown of the components and the underlying mechanics.

### 1C. Lineage Labels (l_i_)

Every individual *i* belongs to one of two lineages, denoted by the label *l*_*i*_:

- **s (Safe Resident):** The established, native population.
- *u* (Unsafe Invader): The invading population trying to colonize the system.

### 2C. The Reproductive Rate Equation (λ_i_)

An individual’s birth rate **λ**_**i**_ is determined by its specific strategy angle *θ*_*i*_ and its lineage *l*_*i*_:

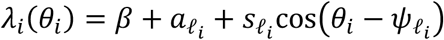

- **β (Baseline Fitness):** The baseline reproductive rate shared by all individuals.
- 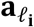 **(Intrinsic Advantage):** A baseline bonus or penalty unique to the lineage (*a*_*s*_ vs *a*_*u*_). For example, if *a*_*u*_ > *a*_*s*_, the invaders have an inherent biological edge.
- 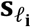 **(Directional Selection Strength):** How heavily the population is penalized or rewarded for deviating from their optimal strategy angle.
- **θ**_**i**_ **(Individual Trait Angle):** The current “intelligence direction” or continuous behavioral state of individual *i*.
- 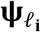 **(Preferred Direction):** The optimal angle for that lineage. The model sets:
- *ψ*_*s*_ = 0 (Safe residents reproduce best when *θ*_*i*_ is close to 0).
- *ψ*_*u*_ = *π* (Unsafe invaders reproduce best when *θ*_*i*_ is close to *π*, completely opposite to the residents).

The cosine term 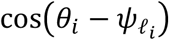 acts as a fitness landscape. Fitness is maximized when an individual’s angle matches its lineage preference (cos(0) = 1) and minimized when it is perfectly misaligned (cos*π* = −1).

### 3C. Inheritance and Death Mechanics

- **Strict Inheritance:** When an individual reproduces, the offspring perfectly inherits both the parent’s exact phenotypic angle *θ*_*i*_ and their lineage label *l*_*i*_.
- State-Neutral Death: An individual’s chance of dying does not depend on its trait angle *θ*_*i*_ or lineage *l*_*i*_. Death is dictated entirely by density regulation (e.g., carrying capacity or overall population crowding).

### 4C. Separation of Learning and Selection

The text highlights a crucial conceptual boundary: “fitness does not directly move an individual’s angle.”

- This means individuals do not physically change or adapt their own *θ*_*i*_ during their lifetime to get more rewards (which would be an active learning or behavior-shifting dynamic).
- Instead, change happens at the population level via natural selection. Individuals with “better” angles simply have more babies, causing those advantageous angles and lineage labels to naturally multiply and dominate the gene pool over generations.

### 5C. Invasion Quantifiers (q_u_)

The model tracks the success of the invasion using the unsafe-lineage frequency *q*_*u*_(*t*), which is the proportion of invaders in the total population:

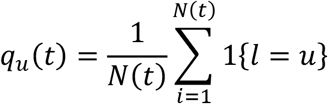

(Where 1{*l*_*i*_ = *u*} equals 1 if the individual is an invader, and 0 otherwise).The invasion simulations follow these operational rules:

- **Initial Setup:** The experiment starts at time *t* = 0 with a tiny invader frequency of 2% (*q*_*u*_(0) = 0.02) dropped into a synchronized resident population.
- **Establishment:** The invasion is deemed successful if, at a designated end time *T*, the invaders make up more than 50% of the population (*q*_*u*_(*T*) > 0.5).
- **Loss:** The invasion fails completely if the invaders drop to 0% (*q*_*u*_(*T*) = 0) by time *T*.

## Appendix D. Numerical validation and reporting framework

The numerical experiments were organized cumulatively so that each additional mechanism was tested against the preceding validated layer.

### Diffusion baseline

For fixed population size, the Euler–Maruyama update is

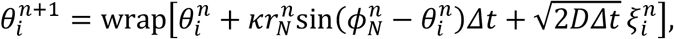

where 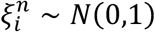 independently. Validation comprises increasing-*N* convergence to *r*_∗_, multiple initial distributions, a below-critical control, and a half-step sensitivity comparison.

### Neutral demographic turnover

Conditional on the current population, births and deaths are sampled by the tau-leap approximation stated in Methods. Parents and deaths are sampled uniformly, so the demographic mechanism is neutral with respect to angle and lineage. Validation includes initial sizes below, at, and above *K*; a carrying-capacity sweep; extinction monitoring; and a half-step comparison.

### Directional selection and lineage invasion

Birth rates are evaluated from the lineage- and angle-dependent fitness function in Section 2.4. Newborns inherit the parent’s angle and lineage. The directional-selection comparison changes the preferred direction while holding the remaining design fixed. Invasion runs begin from *q*_*u*_(0) = 0.02 and vary the unsafe lineage’s intrinsic advantage. Establishment is the finite-time event *q*_*u*_(*T*) > 0.5; loss is *q*_*u*_(*T*) = 0.

### Monte Carlo summaries

If *X* of *M* independent replicates establish, the reported estimate is 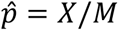. Wilson intervals are used for binomial uncertainty. Convergence slopes are obtained by ordinary least squares after log transformation,

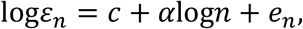

where *n* denotes *N* or *K* and *ε*_*n*_ is the mean absolute deviation from the analytic order-parameter target. These slopes describe the simulated range only.

### Information required for reproducibility

Before submission, the public code archive and manuscript must report the complete parameter table (including *D, κ, β, δ, K, s*_*ℓ*_, *a*_*ℓ*_, *T, Δt*, tail-averaging window, replicate counts, and random seeds) for every figure and table. This information should be generated directly from the executed notebook to prevent transcription error.

